# Distinct distributed brain networks dissociate self-generated mental states

**DOI:** 10.1101/2025.02.27.640604

**Authors:** Nathan L. Anderson, Joseph J. Salvo, Jonathan Smallwood, Rodrigo M. Braga

**Author notes:** corresponding authors: Nathan L. Anderson Rodrigo M. Braga.

## Abstract

Human cognition relies on two modes: a perceptually-coupled mode where mental states are driven by sensory input and a perceptually-decoupled mode featuring self-generated mental content. Past work suggests that imagined states are supported by the reinstatement of activity in sensory cortex, but transmodal systems within the canonical default network are also implicated in mind-wandering, recollection, and imagining the future. We identified brain systems supporting self-generated states using precision fMRI. Participants imagined different scenarios in the scanner, then rated their mental states on several properties using multi-dimensional experience sampling. We found that thinking involving scenes evoked activity within or near the default network, while imagining speech evoked activity within or near the language network. Imagining-related regions overlapped with activity evoked by viewing scenes or listening to speech, respectively; however, this overlap was predominantly within transmodal association networks, rather than adjacent unimodal sensory networks. The results suggest that different association networks support imagined states that are high in visual or auditory vividness.

**Teaser:** Different large-scale brain networks support imagining of visual and audiolinguistic mental content.

## Introduction

Human cognition extends beyond immediate action, encompassing the ability to use imagined mental states to reflect on past experiences and possible future events, both of which support adaptive decision-making. Studies have shown that these self-generated imagined states are often multivariate in nature and rely on both the capacity to imagine scenes with vivid details (Wang et al., 2020; Konu et al., 2020) and to structure thoughts through internal representations of language and speech (Vatansever et al., 2017; Sulfaro et al., 2024; Alderson-Day & Fernyhough, 2015; Carruthers, 2018). Although these mental states are well documented in lab settings (Konu et al., 2021) and in daily life (Mulholland et al., 2023, 2024) we know relatively little about the mechanism through which they emerge.

A key component of self-generated mental states is that they are often accompanied by *mental imagery*—subjective experiences that resemble sensory perception but lack an immediate sensory cause (Kosslyn et al., 2006; Alderson-Day & Fernyhough, 2015; Fernyhough & Borghi, 2023; Sulfaro et al., 2024). For example, for most people (Zeman, 2024), recalling a landmark like Chicago’s Cloud Gate might evoke a mental impression (i.e., visual mental imagery) that is similar to actually seeing the monument. Research has implicated that mental imagery involves the reinstatement of activity within sensory areas that are engaged during perception (Kosslyn et al., 1997; Ganis et al., 2004). For example, O’Craven and Kanwisher (2000) demonstrated that imagining faces or scenes activates regions of ventral temporal cortex that overlap with those activated during visual perception of faces or scenes, respectively. This alignment between imagery and perception is thought to extend across the visual hierarchy (Winlove et al., 2018; Cichy et al., 2012). For example, imagined content can be decoded from primary visual cortex (Naselaris et al., 2015; see also Cabbai et al., 2024). Similarly, when participants are asked to focus on visual details or imagine objects in particular orientations or visual field locations—i.e., features that are encoded in early visual cortex—activity in early visual areas is more reliably evoked (Dijkstra, 2024; Kosslyn & Thompson, 2003; Slotnik et al., 2005; Lambert et al., 2002; see also Chen et al., 1998; Cui et al., 2007). These studies suggest that neural reinstatement of sensory areas is a key correlate of imagined states.

In contrast, studies that examined more complex self-generated states have highlighted that both the processes of mental time travel (i.e., thinking about the past or future) and language-use rely on networks in association cortex rather than sensory cortex. The canonical default network (DN), a distributed brain network involving regions in the posteromedial cortex (e.g., posterior cingulate and retrosplenial cortex), posterior inferior parietal lobe, lateral temporal cortex, and ventral and dorsal medial prefrontal cortices (Buckner et al., 2008) has been repeatedly implicated in mental time travel (Schacter et al., 2007). The DN exhibits heightened activity when individuals are disengaged from the external environment (Shulman et al., 1997)—a context in which they often report thinking about the past and future (e.g., Smallwood et al., 2009; Schacter et al., 2007), or when they actively engage in introspective tasks, such as imagining future scenarios, recollecting autobiographical memories (Spreng & Grady, 2010; Schacter et al., 2007), or making social inferences (Saxe, 2006), whether relevant (Andrews-Hanna et al., 2010) or irrelevant (e.g., Konu et al., 2020) to the task at hand. This broad engagement across diverse self-generated mental states led to the suggestion that the DN plays a broad role in imaginative states, potentially through mechanisms like self-projection (Buckner & Carroll, 2007; Carroll, 2019) or the maintenance of the narrative self (Menon, 2023). However, research has shown that when thoughts include verbal content, increased activation is observed within brain regions that process heard or read sentences (e.g. Amit et al., 2017; Vatansever et al., 2017; Tian et al., 2016; Lu et al., 2023; McGuire et al., 1996; Shergill et al., 2002; Aleman et al., 2005), a set of regions referred to here as the language network or LANG (Fedorenko et al., 2024). Similar to the DN, the LANG network is also distributed across multiple association regions - including inferior frontal, lateral temporal, and posterior middle frontal cortices, as well as the supplementary motor area and potentially further regions (Fedorenko et al., 2024; Braga et al., 2020) - and can be defined using functional connectivity analyses (Braga et al., 2020; Glasser et al., 2016; Lee et al., 2012; Hacker et al., 2013). These studies imply that more complex, integrated self-generated states that include thinking about events or language are often accompanied by activity in association areas of the brain, rather than sensory areas.

It is unclear how the sensory reinstatement hypothesis is reconciled with this role of association networks (i.e., the DN and LANG networks) in supporting self-generated mental states. Emerging evidence suggests the need for refinement in both of these camps. First, recent within-individual functional imaging studies have revealed that the canonical DN comprises at least two parallel distributed networks: DN-A and DN-B (Braga & Buckner, 2017; Braga et al., 2019; see also Andrews-Hanna et al., 2010; Spreng & Andrews-Hanna, 2015; Steel et al., 2021; 2023; Deen & Freiwald, 2021). Activity within these networks is linked to different forms of self-generated thought: DN-A supports ‘mental scene construction’ (Schacter & Addis, 2009) as used when thinking about events in the past or future, while in contrast, DN-B is engaged during social inference-making, such as when considering others’ perspectives or making judgments about social scenarios (DiNicola et al., 2020; 2023; Edmonds et al., 2024; see also Peer et al., 2015; Silson et al., 2019a). These findings suggest some degree of domain-specificity within the canonical DN, and that the appearance of a domain-general role of the DN in self-generated thought may be a consequence of averaging across closely juxtaposed networks.

Second, there has been refinement of the idea that imagery and perception rely on the same brain regions. The initial report by O’Craven & Kanwisher (2000) emphasized that the regions serving imagery and perception of faces and scenes were only partially overlapping. Similarly, it has recently been emphasized that regions involved in perceiving novel scenes or remembering familiar places are actually adjacent, and only partially overlapping (Steel et al., 2021; 2023; 2025; Silson et al., 2019a; 2019b). More detailed analysis, such as that achievable by studying the individual brain (Laumann et al., 2015; Braga & Buckner, 2017; Gordon et al., 2017), may be necessary to conclusively resolve the underlying functional organization of these regions. Interestingly, the adjacent ‘scene perception’ and ‘place memory’ regions are positioned in a way that respects the unimodal-transmodal gradient of brain organization: scene perception areas are located in more posterior locations of posteromedial and lateral parietal cortex, closer to unimodal visual cortex, whereas the place memory areas tend to be more anterior, closer to transmodal association cortex (Silson et al., 2019a; 2019b; Bainbridge et al., 2021; Steel et al., 2021; 2023; 2025). Silson et al. (2019a) further suggested that the boundaries of the canonical DN actually bisect these adjacent regions, with place memory areas within the DN, and scene perception areas outside. These findings outline a more nuanced model and raise three putative principles, whereby regions supporting external (perceptual) and internal (self-generated) representations of similar content are (i) adjacent or partially overlapping, (ii) positioned along the unimodal-transmodal gradient, and (iii) potentially differentiated by their underlying connectivity structure with the rest of the brain (i.e., the networks in which they belong).

Do these principles extend to other forms of mental imagery? Research on non-scene-related forms of imagery supports interesting parallels. Partial overlap has been noted between perception and imagery for sounds and music (Halpern & Zatorre, 1999; Herholtz et al., 2012; Kraemer et al., 2005) as well as imagining speech (e.g., Tian et al., 2016; Lu et al., 2023; Shergill et al., 2002). Studies diverge on whether the recruitment of primary and secondary sensory areas occurs during auditory imagery, suggesting that potentially the extent of the auditory sensory hierarchy might be reinstated depending on the imagined content (Yoo et al., 2001; Bunzeck et al., 2005; Tian et al., 2016; Shergill et al., 2001). Further, a similar unimodal-transmodal topographic organization may be present: Herholtz et al. (2012) provided maps showing adjacent or partially overlapping regions, with music imagery regions encircling auditory cortical regions activated by listening to music. The juxtaposed and partially overlapping regions were positioned such that imagery-related regions were closer to transmodal association cortex than perceptual regions were (see also Kleber et al., 2007; Spagna et al., 2021). Thus, along many dimensions the relationship between auditory imagined and perceptual states may recapitulate observations from the scene imagery literature, but in a different sensory modality and set of brain regions.

The above studies underscore a major difficulty in studying self-generated mental states: the specifics of what participants think about can have a substantial effect on the activation patterns. Conventional analysis strategies, which assume that participants are engaging with the task as intended, might be improved upon by accounting for trial-wise variation in thought content (Yarkoni et al., 2009; Smallwood et al., 2021; DiNicola et al., 2023). Recently, multi-dimensional experience sampling (mDES) has been used to highlight (i) the neural systems engaged by states that emerge from the processing of perceptual input (McKeown et al., 2023) and movies (Wallace et al., 2025), (ii) self-generated states such as oN-task thought (Konu et al., 2020; Turnbull et al., 2019; Iwata et al., 2024) and (iii) states of task-evoked imagination (Villena-Gonzalez et al., 2018; Zhang et al., 2022). Demonstrating the potential benefits of accounting for trialwise variance in mental states, a recent mDES study showed that trials in which participants reported thinking more about scenes corresponded to increased activity within the full set of distributed regions of DN-A (DiNicola et al., 2023; see also Gilmore et al., 2021), including but extending beyond the posteromedial and lateral parietal “place memory” areas (Steel et al., 2021, 2023; Silson et al., 2019b). This activity pattern suggests that entire transmodal networks may be recruited during the evocation of mental imagery, rather than only regions bordering perceptual regions. The increased sensitivity of accounting for trial-wise variation in thought-content might yield similar insights into other forms of self-generated states.

Here, we investigated whether similar principles were conserved across different forms of self-generated thought. Participants were asked to imagine several prompts in the scanner, and then rate their mental content on several experiential features using mDES (Smallwood et al., 2016; Smallwood et al., 2021). We tested how brain activity during these two types of imagined states overlapped with activity during the perception of related content. Finally, we asked how this functional organization relates to within-individual maps of transmodal distributed networks and unimodal sensory networks.

## Materials & Methods

### Overview

#### Subjects and sessions

Ten adults (6 female, ages 22-36, mean age 26.6, 9 right-handed) from the local community were recruited as part of the ‘Detailed Brain Network Organization’ (DBNO) study. Details regarding this dataset have been previously described in Kwon et al. (2024), Edmonds et al. (2024), and Salvo, Anderson et al. (2024). Participants had normal hearing and normal or corrected to normal vision, and no history of neurological illness. All participants provided written consent to take part in the study and were compensated for participation.

Procedures were approved by Northwestern University’s Institutional Review Board. Participants were invited to 8 magnetic resonance imaging (MRI) sessions, each of which included several cognitive tasks including a passive fixation (“resting-state”) task for network mapping using functional connectivity (FC). Prior to the first MRI session, participants were trained on all the tasks and were extensively coached about strategies for staying still in the scanner to improve data quality. Participants’ heads were padded using inflatable cushions to restrict head motion. Participants were informed that the initial 1-2 MRI sessions would serve as a trial to assess compliance and whether they wanted to continue participation. Based on these criteria, 2 subjects were not invited back for additional sessions, leaving 8 subjects who completed all 8 MRI sessions (4 female, ages 22-36, mean age 26.75, 7 right-handed). This led to a total of 60.8 hours of fMRI data collected, including 7.6 hours per participant, for precision functional mapping.

#### Passive fixation task

To allow estimation of large-scale networks within each individual using FC, participants completed a passive fixation task (REST) in the scanner. Participants were shown a crosshair in the center of the screen and instructed to fixate on the crosshair for the duration of the task, keeping their eyes open and blinking normally. The task lasted ∼7 minutes, and each participant completed two runs in each session (total: 16 runs, ∼112 minutes per participant). One participant (S2) completed an additional REST run to replace a poor-quality run that had been excluded in a prior session (see *MRI quality control*). Two runs of REST were collected in each MRI session, and these were the first and last runs in each session.

#### IMAGINE Task Overview and Stimuli

Participants took part in a cued imagining task (IMAGINE) in the scanner, in which they were asked to read prompts and imagine the prompted content, then press buttons to report the auditory and visual vividness of the imagined content. After the scan, participants retrospectively answered multiple questions about their thought content in the scanner for each trial (mDES). Participants were given detailed instructions about the task (see script included in Supplementary Materials) and practice trials so they were familiar with the task structure prior to scanning. Instructions included the following descriptions for vividness rating levels: “‘*Nothing’* means you did not experience any mental images or sounds at all, ‘*Vague*’ means you think you did experience mental images or sounds, but they were dim or dull or not very clear, ‘*Moderate*’ means you did experience mental images or sounds, and you feel that they were relatively clear or vivid, ‘*Vivid*’ means you did experience mental images or sounds, and they were very clear and vivid, almost similar to if you were actually hearing or seeing the item in real life” (see Marks, 1973). Participants were reminded of instructions before each MRI session and were given a chance to practice the task outside of the scanner.

Two hundred sentences were created as prompts for the IMAGINE task. Candidate sentences were generated by authors NLA and RMB and checked by others to eliminate sentences that were difficult to interpret or produced unintended associations. Forty prompts were created targeting five categories: Scenes, Faces, Perceptual Speech (i.e., imagining listening to someone else speak), Inner Speech (i.e., imagining specific words using an “internal voice” or “inner monologue”), and Sounds. Each prompt began with the word “Imagine…” followed by instructions, to create prompts such as “Imagine an art studio” or “Imagine hearing a couple arguing”. The full list of prompts is shown in Table S1. The average prompt length was 32.4 characters (sd = 5.4, range = 19-45) and prompt length was matched across conditions (mean for Scenes: 32.0 characters; Faces: 32.3; Perceptual Speech: 32.4; Inner Speech: 32.5; Sounds: 32.7). Prompts were selected to evoke high visual vividness for the scene- and face-imagining trials, and high auditory vividness for the sound and speech conditions.

#### In-scanner IMAGINE task

Each of the 8 MRI sessions included an fMRI run of the IMAGINE task lasting 614 seconds (approx. 10.2 minutes), leading to a total of 4,912 seconds (approx. 80.9 minutes) of IMAGINE task data collected per person prior to quality control. Each run included 25 prompts (one per trial), with five trials per category. Prompt order was counterbalanced using *optseq2* (Dale, 1999). Participants were presented with a prompt written on the screen and given 7 seconds to read the prompt and covertly imagine the prompted content. They were then asked to rate the visual vividness of what they imagined on a 4-point scale (Nothing – Vague – Moderate – Vivid), followed by a rating of auditory vividness on the same scale (see Figure 1A for schematic). Half of the runs included the visual vividness rating first, and half included the auditory vividness rating first, alternating each session. The rating period lasted 3s for each rating. Participants responded by pressing buttons with the middle and index fingers of each hand (corresponding to increasing vividness ratings moving from left to right). The response button meanings were shown on the screen beneath the vividness questions during the rating period (Fig. 1A). Each trial (consisting of the imagining period followed by the two ratings) was preceded by a jittered 10-11 second inter-trial fixation period that served as a trial-specific baseline. To reduce the time between in-scanner and follow-up experiments, in each MRI session the IMAGINE task was the penultimate task, followed by a 7-minute passive fixation task.

**Fig. 1:**
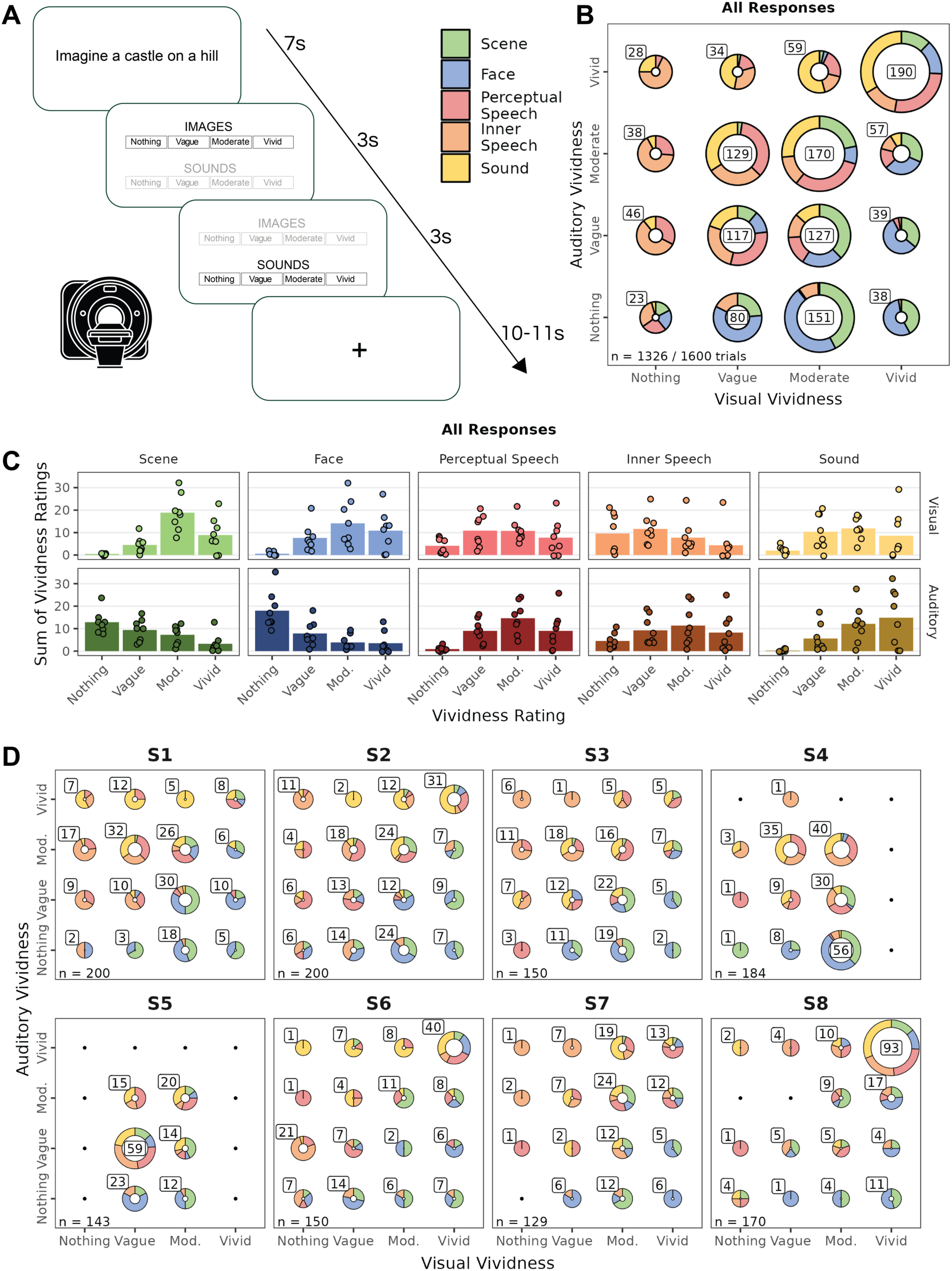
The imagination task successfully modulated the visual and auditory vividness of participants’ mental content in the scanner. A) Schematic of a task trial, wherein participants were cued to imagine diJerent scenarios and then rate the vividness of their imagined content. B) Plot of vividness ratings across all trials from all participants, divided by stimulus condition. Scene and Face trials were generally rated as more visually vivid, while Perceptual Speech, Inner Speech, and Sound trials were rated as more auditorily vivid. C) The same pattern can be seen when trials are separated into visual (top row) and auditory (lower row) vividness ratings, for each stimulus condition. Points represent individual participants, while bars represent the average. D) Within-individual plots of vividness ratings, similar to panel B, show that the separation of visual and auditory vividness across conditions held in each participant, though there were appreciable inter-subject diJerences.

#### Out-of-scanner IMAGINE follow-up task

Due to practical limitations, such as the amount of time available in the scanner, only the two vividness ratings were collected during each trial of the in-scanner task. To comprehensively characterize participants’ thought-content, an additional set of questions or ‘thought-probes’ were collected in a follow-up experiment. Fifteen thought-probes were created to explore different facets of imagining, including content (e.g., “I envisioned the location of objects, people or places”), temporal dynamics (e.g. “I imagined a sequence of events unfolding in my mind”), imagery source (“To imagine this, I drew on a memory of a specific fact, place, person, or object”), or performance (e.g. “I found it difficult to imagine this item”). The full list of thought-probes is shown in Table 1. The thought-probes were chosen to differentiate potential functions of the DN and LANG networks. For example, DN-A has been linked with episodic projection (DiNicola et al., 2023), as captured by the thought-probe “I thought about an event from my past or in the future”, while LANG is associated with processing language, hence the thought-probe “My thoughts included specific, identifiable words or phrases”. Similar probes have been used effectively in other task paradigms to identify brain correlates and capture individual differences in experience (see Wallace et al., 2025; Turnbull et al., 2019).

**Table 1:** Multi-dimensional experience sampling (mDES) survey questions collected during the out-of-scanner follow-up task. Participants answered 17 questions about each scenario imagined during that day’s scanning session. Ratings for visual vividness and auditory vividness were presented identically to the in-scanner question. Subsequent questions targeted features of the imagined content such as language use, memory specificity, task performance, and other domains. Participants responded using a 1-9 scale from Strongly Disagree to Strongly Agree. The Focused and Ease probes were reverse-coded for analyses.

| Label | Follow-up Prompt | Rating Scale |
| --- | --- | --- |
| Visual Vividness* | How vividly did you imagine mental images? | Nothing - Vague - Moderate - Vivid<br>(4-point scale) |
| Auditory Vividness* | How vividly did you imagine mental sounds? |  |
| Locations | I envisioned the location of objects, people, or places | Strongly disagree - Neutral - Strongly agree<br>(9-point scale) |
| Immersion | I felt immersed in my thoughts, as if I was experiencing the item(s) in real life |  |
| Sequence | I imagined a sequence of events unfolding in my mind |  |
| Elaboration | My thoughts included elaboration and detail beyond what the prompt included |  |
| Speech or Language | My thoughts involved speech or written language |  |
| Internal Voice | My thoughts included my internal voice or 'inner speech' |  |
| Specific Words/Phrases | My thoughts included specific, identifiable words or phrases |  |
| Other People | I thought about other people |  |
| Past/Future Event | I thought about an event from my past or in the future |  |
| Specific Memory | To imagine this, I drew on a memory of a specific fact, place, person, or object |  |
| Feelings/Preferences | I thought about feelings, preferences, or emotions |  |
| Abstract Concepts | I thought about abstract concepts, such as truth, beauty, or fairness |  |
| Success | I was able to imagine what was asked successfully |  |
| Focused** | My mind wandered and I was not focused when imagining this |  |
| Ease** | I found it difficult to imagine this item |  |

\*Also administered in-scanner
\*\*Reverse-coded for analysis

Immediately after each MRI session, prior to changing out of the hospital scrubs, participants were taken to an interview room to perform the follow-up task. On average, 3.2 minutes elapsed between the participant exiting the scanner and beginning the computerized task (range: 1-7 min). Participants were re-presented with the 25 prompts seen in the scanner, one at a time, and asked to respond to 17 thought-probes on a series of scales (see Fig. 2A for schematic). Participants were instructed to respond regarding what they had imagined for that prompt while in the scanner. Participants first provided the visual and auditory vividness ratings again using the same 4-point scale to assess test-retest reliability, and then completed the additional 15 thought-probes on a 9-point scale (from Strongly Disagree to Strongly Agree; see Table 1). Data were collected and stored using REDCap electronic data capture tools hosted at Northwestern University (Harris et al., 2009; Harris et al., 2019). Participants took an average of 21.5 minutes to complete the follow-up task (range: 12-47 min).

**Fig. 2:**
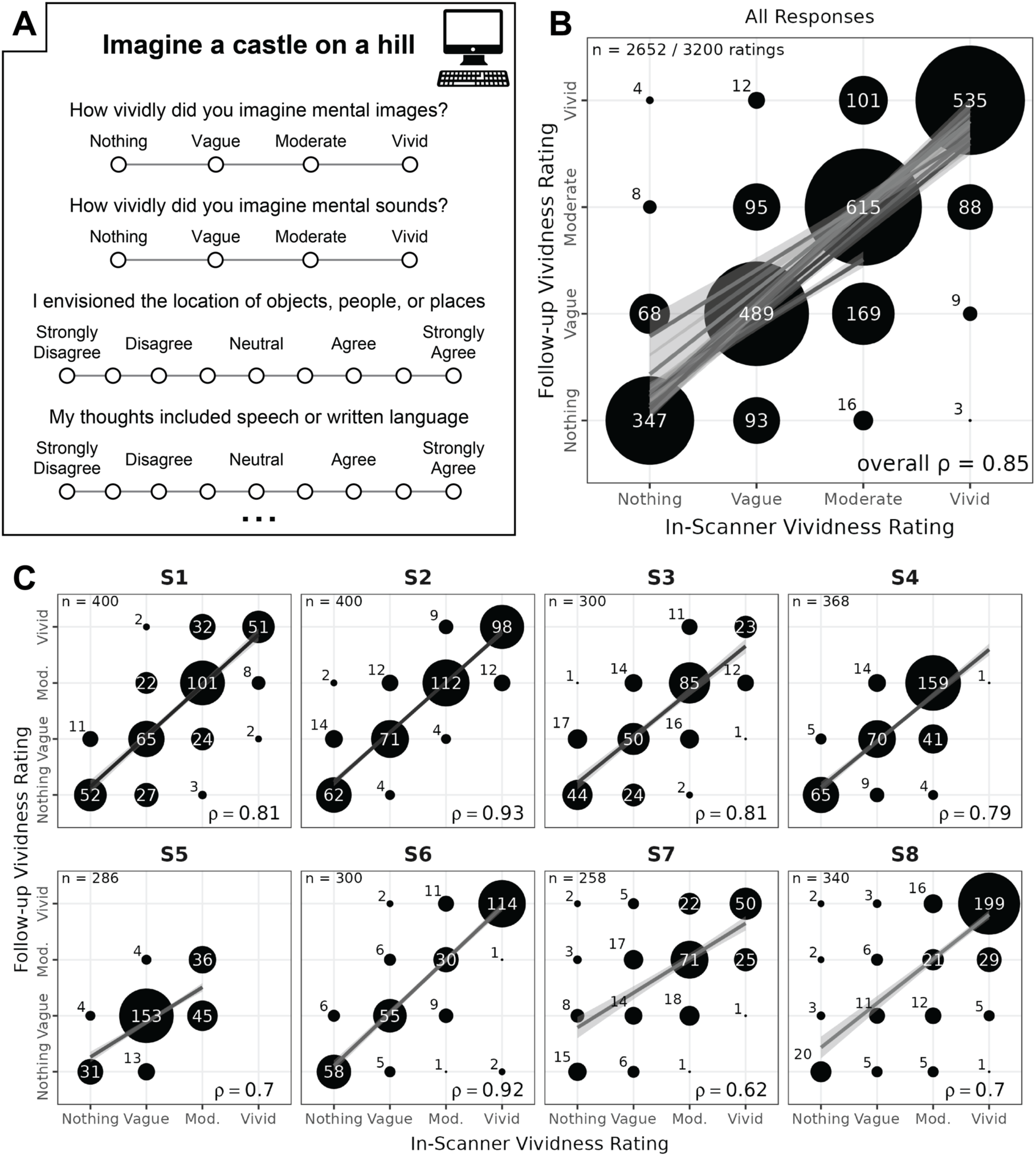
Post-scan follow-up vividness ratings showed strong correspondence to in-scanner vividness ratings. A) Schematic of out-of-scanner survey. Participants completed an out-of-scanner questionnaire with follow-up questions (i.e., multi-dimensional experience sampling; mDES) where they were asked questions about the features of their imagined content during the in-scanner task. Questions were rated on a 1-4 scale (vividness) or 1-9 agreement scale (all others; see full list of follow-up questions in Table 1). B) Confirming that the follow-up responses corresponded to participants’ in-scanner experiences, vividness ratings inside and outside the scanner showed high correlation (overall Spearman’s rho = 0.85) across all subjects. C) This same pattern held in each participant, with correlation remaining high (Spearman’s rho ranging from 0.62 to 0.93).

#### Visual category (VISCAT) localizer

To allow definition of brain regions related to perception of visual images and language within each individual, participants also completed two perceptual localizer tasks in each MRI session (VISCAT, SPEECHLOC). The first was a 4-minute one-back task featuring images of different visual categories (VISCAT; see Saygin et al., 2016). Participants were shown square grayscale images that depicted a scene, a face, a pseudoword, an object, or a visually scrambled image (taken from the scene, face, or object categories). Pseudowords were generated using the Australian Research Council (ARC) Nonword Database (Rastle et al., 2002), while the other four categories were taken from databases provided by Talia Konkle (Konkle & Oliva, 2011). Participants were asked to press their left index finger button if the image was novel, and their right if the image matched the previous image. Each image was displayed for 0.7 seconds, with a 1.3 second interstimulus interval when a fixation crosshair appeared on the screen. Images from each category were shown sequentially in a block design, with 2 blocks per category. There were 9 images in each block. The task consisted of a total of 10 blocks, which included 75 unique images and 15 repeated images. Five blocks of different categories were presented sequentially, and an 18-second fixation period was presented before, in between, and after the 5-block groups that served as a baseline. Categories were counterbalanced within (e.g., all categories were presented in the first 5-block group, and then the order was reversed in the second group) and across runs to minimize order effects.

#### Auditory language (SPEECHLOC) localizer

The second localizer task was a 7-minute auditory language localizer (SPEECHLOC; based on Scott et al., 2017), where participants listened to audio clips of clear, unfiltered speech and distorted, unintelligible speech. Audio clips were taken from recordings of TED talks, The Moth radio hour, and Librivox audiobooks. Two clips were taken from each recording, with one clip left intact and the second distorted by filtering (this filtering process is described in detail in Salvo, Anderson et al. 2024). This was done to provide a contrast condition that included sound clips which preserved certain sound characteristics (speaker intonation, speed, etc.) but did not contain comprehensible speech.

Audio during the task was presented through Sensimetrics S14 MRI-compatible earphones (Sensimetrics, Gloucester, MA, USA). To ensure that the clips were audible over the scanner noise, at the start of each MRI session, prior to data collection, participants were played two speech clips (one clear, one distorted) in the scanner while the same MR sequence was running. Participants used the provided button box to adjust the volume themselves to a comfortable level where the speech could be clearly heard and were asked to describe to the experimenters the topic of the clearly-presented story. Each audio clip lasted 18 seconds and was followed by a tone lasting 0.15 seconds. Following each tone, participants had 2 seconds to press a button with their right index finger to confirm they were paying attention. A fixation crosshair appeared on the screen throughout the run and participants were asked to maintain fixation. All other cues and stimuli were auditory. Each task run included 8 clear and 8 distorted 18s audio clips, which were interleaved during task presentation. Each run included four blocks, with four clips presented in each block. Each block included two clear clips and two distorted clips, with a 14-second fixation cross (‘+’) presented between blocks. Within each block, clips were presented in a counterbalanced order, e.g., CDDC+DCDC+CDCD+DCCD (where “C” is a clear clip and “D” is a distorted clip, and + is fixation), ensuring an even distribution of conditions across the run. Block order was also counterbalanced across runs.

In each session, participants also completed other tasks (e.g., targeting social cognition) that are not described in this report (see Edmonds et al., 2024; Kwon et al., 2024; Salvo, Anderson et al., 2024).

#### MRI data acquisition

Data were collected on a Siemens 3.0 T Prisma scanner (Siemens, Erlangen, Germany) at the Center for Translational Imaging at Northwestern University’s Feinberg School of Medicine in Chicago. Two anatomical images were collected: a T1-weighted image in the first MRI session (TR = 2,100ms, TE = 2.9ms, FOV = 256 mm, flip angle = 8°, slice thickness = 1 mm, 176 sagittal slices parallel to the AC-PC line) and a T2-weighted scan in the second session (TR = 3,000ms, TE = 565ms, FOV = 224 mm x 256 mm, flip angle = 120°, slice thickness = 1 mm). Both scans included volumetric navigators (Tisdall et al., 2012) from the Adolescent Brain Cognitive Development study (Hagler et al., 2019). Functional MRI data were collected using a 64-channel head coil with a multi-band, multi-echo sequence with the following parameters: TR = 1,355ms, TE = 12.80ms, 32.39ms, 51.98ms, 71.57ms, 91.16ms, flip angle. = 64°, voxel size = 2.4mm, FOV = 216 mm x 216 mm, slice thickness = 2.4 mm, multiband slice acceleration factor = 6 (see Poser et al., 2006; Lynch et al., 2021).

#### Quality control

Head motion was estimated using FSL’s *MCFLIRT* (Jenkinson et al., 2002). Runs were automatically excluded from analysis if motion exceeded predetermined thresholds of framewise displacement (FD) > 0.4mm or absolute displacement (AD) > 2.0mm. Runs were flagged for visual inspection if motion exceeded more stringent thresholds (FD > 0.2mm, AD > 1.0mm), and the whole run was then excluded if motion could be clearly seen in the raw data. On this basis, each subject retained at least 6 runs of the IMAGINE task (S1: 8; S2: 8; S3: 6; S4: 8; S5: 6; S6: 6; S7: 6; S8: 7 runs) leaving an average of 70.1 minutes of data per participant (range:61.2-81.6 min). Additional quality control was performed based on the behavioral performance, where we excluded individual trials in which the participant did not respond to both vividness ratings (i.e., missed responses; 36 trials excluded; S1: 0; S2: 0; S3: 0; S4: 12; S5: 7; S6: 0; S7: 12; S8: 5) or pressed multiple buttons (13 trials excluded; S1: 0; S2: 0; S3: 0; S4: 4; S5: 0; S6: 0; S7: 9; S8: 0). After all quality control measures, each subject provided at least 129 trials from the IMAGINE task (range: 129-200).

For the REST task, runs were excluded based on the same motion criteria described above. After quality control, each participant retained at least 10 runs (range: 10-16). For all participants except one (S5), half of the data was set aside for replication analyses (not used in this study), leaving 49-60 minutes of data for each participant (S1: 56 min; S2: 56; S3: 56; S4: 56; S5: 60; S6: 49; S7: 49; S8: 49).

Runs for the remaining two localizer tasks were quality controlled for motion using the same procedure as for IMAGINE and REST. Additionally, runs were excluded in their entirety based on poor performance using behavioral responses. For the one-back VISCAT, runs with an error rate above 20% were excluded. For SPEECHLOC, runs with more than 20% missing button-presses were excluded. After quality control, all 8 runs were retained for each subject for VISCAT. For SPEECHLOC, at least 7 runs were retained per subject (S1: 8; S2: 8; S3: 8; S4: 7; S5: 7; S6: 8; S7: 7; S8: 8).

#### MRI data preprocessing

Blood-oxygenation-level-dependent (BOLD) data were preprocessed using a custom processing pipeline “iProc” (Braga et al., 2019) that is optimized for within-individual alignment of data from multiple runs and sessions, and minimizing blurring through reduced smoothing and interpolation. Additional steps were included to account for the multi-echo data (outlined below). Each individual’s data was processed separately. The first 9 volumes (∼12 s) of each run were discarded to remove the T1 attenuation artifact. Next, a mean BOLD template was created as an interim stage for data registration by averaging the first echo of all included runs of all tasks, to reduce bias towards any one particular run. The participants’ T1 anatomical images were used to create a native space template. These templates were used to build four matrix transforms to align each BOLD volume 1) to the middle volume of the same run for motion correction, 2) to the mean BOLD template for cross-run alignment, and 3) to the native space template, and 4) to the Montreal Neurological Institute (MNI) International Consortium for Brain Mapping (ICBM) 152 1-mm atlas (Mazziotta et al., 1995). The four transforms were composed into two matrices which were then applied to the original volumes to register all volumes to the T1 anatomical template (matrices 1-3) and to the MNI atlas (matrices 1-4) in a single step. Visual checks were incorporated into each registration step of the pipeline.

Motion correction transforms were calculated using rigid-body transformation based on the first echo, which has less signal dropout and better preserves the shape of the brain. Registration matrices were calculated based on the first echo, and then applied to all the echoes. Once data were projected to the T1 native space, the five echoes were combined to approximate local T2* by weighting each echo according to its temporal signal-to-noise (tSNR) ratio and echo time (Heunis et al., 2021). Briefly, the tSNR is calculated for each echo, which is then weighted by the echo time. This weighted tSNR is then divided by the sum of all weighted tSNRs (i.e., for all echoes). The resulting image is multiplied by the echo’s original intensity image, and these are summed across all echoes to create the final optimally combined image.

#### Functional connectivity preprocessing

Nuisance variables were calculated for each run, including average signal from deep white matter and cerebrospinal fluid using masks hand-drawn in MNI space that were back-projected to the native space. Additional nuisance variables included the 6 motion parameters and whole-brain (global) signal, and their temporal derivatives. Data were then bandpass filtered using a range of 0.01–0.1 Hz using *3dBandpass* (AFNI v2016.09.04.1341; Cox, 1996, 2012). Data were then projected to a standard cortical surface (fsaverage6, 40,962 vertices per hemisphere; Fischl et al., 1999), and smoothed with a 2mm full-width at half-maximum kernel along the surface. This kernel size was chosen based on prior work (Braga & Buckner, 2017; Braga et al., 2019) to retain functional anatomic detail while limiting noise (i.e., speckling) in the functional connectivity maps.

#### Network estimation

Prior to any analysis of the task data, the REST data was used to create subject-specific estimates of large-scale networks using functional connectivity following our previously used procedures (Braga & Buckner, 2017; Braga et al., 2020). First, functional connectivity matrices were calculated for each REST run by computing vertex-vertex Pearson’s product-moment correlations. The matrices were then z normalized using the Fisher transform, averaged across all runs within each individual, then converted back to r values using the inverse Fisher transform. These within-individual average matrices were used to perform a seed-based analysis. Seeds were manually selected in the left lateral prefrontal cortex to delineate 7 networks (Default Network A [DN-A], Default Network B, Language Network [LANG], Frontoparietal Network A, Frontoparietal Network B, Dorsal Attention Network A, and Dorsal Attention Network B [referred to here as dATN]; see Braga & Buckner, 2017; Braga et al., 2020; Edmonds et al., 2024; Kwon et al., 2024). Analyses here focused primarily on DN-A and LANG. We then used a multi-session hierarchical Bayesian Model (MS-HBM) approach to define networks using a data driven algorithm applied to the same data (Kong et al., 2019). This method provides individual-specific network estimates by integrating priors from multiple levels (e.g., group atlas, cross-individual and cross-run variation) to stabilize network estimates. For each individual, we generated clustering solutions with k values (i.e., number of clusters) between 12–18, and selected the lowest level of clustering which separated the networks of interest as defined by the seed-based approach. Based on these criteria, a 14-network solution was selected. The MS-HBM algorithm also allowed definition of networks beyond the *a priori* selected networks, including those covering auditory cortex (surrounding bilateral Heschl’s gyrus; referred to here as AUD), ventral somatomotor cortex (surrounding bilateral inferior central sulcus; referred to here as SMOT), and peripheral visual cortex (referred to here as VIS-P; see Du et al., 2024). In one subject, S8, MS-HBM identified a LANG network that did not align well with the seed-based analysis or the task activation maps for SPEECHLOC (as discussed in Salvo, Anderson et al., 2024). Therefore, in this subject, we took a higher clustering solution (*k* = 15) which produced a LANG network that better fit the task maps.

#### Task activation maps

All MRI tasks (except REST) were analyzed using FSL’s *FEAT* (Woolrich et al., 2001). Each hemisphere of each surface-projected task run was input into a separate general linear model (GLM). The task was modelled using explanatory variables that were convolved with a double-gamma hemodynamic response function using *FEAT*.

For the IMAGINE task, independent regressors were specified for each of the 25 trials, covering the period from the onset of the prompt to the end of the 7-second imagining period. An additional 25 regressors were specified to cover an equivalent 7-second period taken from the intertrial fixation periods that preceded each trial. Beta values were extracted for each regressor, and we contrasted beta values from imagination vs. fixation periods for each trial using *FEAT*. The fixation periods acted as independent baselines for each trial, ensure ensuring that the estimates of the contrast of parameter estimates were independent for each trial (Hassabis et al., 2014; DiNicola et al., 2020). A separate 25 regressors modelled the response period for each trial.

For VISCAT, regressors were specified that covered the entire block for each of the visual category types (e.g. scenes, faces). Only the scenes condition was of interest in this report, and so the beta values extracted were composed of a contrast between scenes vs. the conjunction of faces, objects, and pseudowords.

For SPEECHLOC, regressors were specified that included all 18-second presented audio clips for each of the two speech types (comprehensible and distorted). Beta values were calculated for the comprehensible and distorted speech, and a contrast of parameter estimates was calculated for each run.

#### Incorporation of trialwise mDES responses

The data from the IMAGINE task were analyzed in three ways. First, we performed a simple ‘condition-level’ contrast between different prompt categories. For the scene-imagining condition, beta values for Sound trials were multiplied by -1, and then the values for all Scene and Sound trials were averaged together (forming a contrast between Scenes and Sounds). For the language map, the same procedure was followed, except the Perceptual Speech and Inner Speech trials were first averaged together and then contrasted with the Sound trials to create a single contrast map. An additional map contrasted trials from the Perceptual Speech category against trials from the Inner Speech category.

For the second analysis, the behavioral ratings provided for each trial during the post-scan multi-dimensional experience sampling (mDES) task were used to group trials based on features reported by the participants themselves, rather than the *a priori* categories (see Varrier & Finn, 2022). To calculate a ‘scene construction map’, a composite score was created for all trials using the sum of the Visual Vividness and Locations thought-probes (based on DiNicola et al., 2023). Beta values from trials that were in the top 20% of ratings on this composite were contrasted against trials in the bottom 20%. If items were tied at the cutoff rating, subsets of these trials were randomly selected to achieve 20% of trials. The ‘language map’ was calculated similarly, except the summed composite was composed of the Auditory Vividness, Speech or Language, and Specific Words/Phrases thought-probes.

For the third analysis, behavioral data for IMAGINE were subjected to two different methods of dimensionality reduction: hierarchical clustering and principal component analysis (PCA). A correlation matrix was created for each probe (including the vividness ratings from the in-scanner task, as well as the 15 additional thought-probes from the out-of-scanner follow-up task) using pair-wise Pearson’s *r*. The matrix was then hierarchically clustered using the *hclust* function from the *stats* package in R (R & *stats* version 4.1.1; R Core Team, 2021), using the *ward.D2* method, with 3 clusters specified (the number of clusters that result when the clustering dendrogram is cut at a length of 0.5). Additionally, the behavioral data were analyzed using a PCA, implemented through the *prcomp* function in R’s *stats* package. The first two principal components each explained more than 10% of the variance in the data and were retained for further analysis. In both reduction approaches, the behavioral data from all subjects after quality control was used. All analyses were repeated creating individual clustering and PCA results for each subject; the results were very similar, so the omnibus approach was chosen.

Correspondence between the two principal components and the trial-wise activation data was then examined (McKeown et al., 2023; Wallace et al., 2025; Konu et al., 2020; Turnbull et al., 2019). A higher-order GLM was created using regressors that weighted trials according to the loadings of each trial onto the first two principal components. This produced vertex-wise beta values representing how each vertex was associated with the trial-wise variance captured in PC1 and PC2. These beta maps were z-scored to create two separate maps.

Finally, overlap between brain activity during perceiving and imagining was determined using relevant maps for each condition (viewing and imagining scenes, listening to and imagining speech). All maps were thresholded at z = 1.5 and then binarized. Regions of overlap were identified, and then compared to the spatial extent of relevant target networks (for scenes: Default Network A [DN-A], dorsal Attention Network [dATN], and peripheral visual cortex [VIS-P]; for speech, Language network [LANG], primary auditory cortex [AUD], and somatomotor cortex [SMOT]). A Dice coefficient was calculated for each network, quantifying the network’s overlap with task activity. Only vertices present in the network and/or overlap regions were included in the calculations.

## Results

### Vividness of visual and auditory imagery was successfully manipulated by the task

As part of the IMAGINE task, participants were prompted to imagine several scenarios in the MRI scanner, and then provide ratings after each scenario regarding the degree of visual and auditory vividness of their imagined content. Although we designed the prompts to elicit visual and auditory imagery, isolating these features can be difficult; prompts designed to evoke auditory imagery (e.g., “Imagine the sound of a waterfall”) might elicit visual imagery, and vice versa. Analysis of in-scanner vividness ratings (Fig. 1A) showed that the prompts were successful in selectively modulating visual and auditory imagery, to varying degrees. A consistent pattern was found across all subjects (Fig. 1B & 1C), where imagined states during the visual categories (Scenes, Faces) were generally rated as higher in visual and lower in auditory vividness (see lower right quadrant in Fig. 1B), whereas trials in the auditory categories (Perceptual Speech, Inner Speech, Sounds) showed the opposite pattern (see upper left quadrant in Fig. 1B). Participants’ vividness ratings therefore aligned with the intended categories, implying that participants were following instructions and responding appropriately, and likely experiencing imagined states with different levels of auditory and visual content.

Analysis of vividness ratings within each individual showed that all subjects (S1-S8) differentiated visual and auditory categories (Fig. 1D), confirming the group-level trend. Some subjects reported lower overall experiences of vivid imagining (e.g., S4, S5) even if their responses still differentiated the categories. This aligns with research showing that vividness levels vary across people (Betts, 1909; Zeman et al., 2015). Supplementary Figure S1 shows each participants’ vividness ratings broken down by category, underscoring that some subjects reported different degrees of separation of auditory or visual imagery across categories, as well as different ranges of responses. Some subjects reported that some trials led to mental sensations that were very vivid in both visual and auditory domains (Fig. 1D; e.g., S6, S8), implying that the prompts often evoked integrated, multisensory experiences. This heterogeneity underscores the need for approaches that can account for trial-wise variation in experienced content.

### Participants’ out-of-scanner responses reflected their in-scanner experiences

Once the MRI session was completed, participants took part in a follow-up mDES experiment (Fig. 2A) where they answered additional questions about the properties of their mental content experienced during the in-scanner IMAGINE task. The visual and auditory vividness ratings were repeated in the mDES experiment, to determine the degree to which participants’ out-of-scanner responses successfully referenced their in-scanner mental states. In- and out-of-scanner vividness ratings were highly correlated (Spearman’s ρ = 0.85; Fig. 2B) and did not differ between visual and auditory ratings (overall visual: ρ = 0.84; overall auditory: ρ = 0.86). Correspondence in each individual participant varied but was also high (range: ρ = 0.62-0.93; Fig. 2C). This consistency provides evidence that participants’ out-of-scanner responses were related to their in-scanner mental states.

Quality control was performed on the mDES responses. Review of participant data showed no consistent instances of stereotyped response patterns (e.g. selecting the same rating for multiple questions in a row, or zig-zagging patterns) that could indicate participants were ignoring instructions. Additionally, because one of the experience sampling probes regarded difficulty (“I found it difficult to imagine this item”), while another regarded success (“I was able to imagine what was asked successfully”), a negative correlation between these probes indicated that participants were responding accurately (range: ρ = - 0.60 to -0.92). One participant (S3) appeared to have interpreted the question about mind-wandering (“My mind wandered and I was not focused when imagining this”) in the opposite direction compared to the other participants (i.e., S3 reported high mind wandering when they also reported successfully imagining the prompt), so their responses were reverse-coded for the mind wandering question to align with other subjects. The full list of experience sampling probes is displayed in Table 1.

### Multi-dimensional sampling (mDES) reveals covariance structure of imagined states

We leveraged the mDES responses to explore how participants’ mental states varied across trials (i.e., prompts). Dimensionality reduction was performed using principal components analysis and hierarchical clustering, seeking converging evidence for structure in the mDES responses.

Fig. 3A shows the pairwise correlation matrix between individual mDES responses across all participants, which highlights that some responses formed clusters (e.g., see pairwise correlations above ρ > 0.5). This was confirmed by the data-driven clustering analyses. The principal components analysis resulted in two principal components that explained more than 10% of the variance in the data (Fig. 3B & 3A, right). The first component (PC1) explained 36.2% of the variance and related positively to all the probes, indicating a relation to general successful performance of the task (i.e., higher PC1 loadings represent higher ratings in general). However, this component also had relatively higher scores for probes such as Visual Vividness, Locations, Immersion, and use of Specific Memories (see purple bars in Fig. 3A, right). We therefore refer to PC1 as representing trials high in ‘Scene Construction’ for brevity in the rest of the text.

**Fig. 3:**
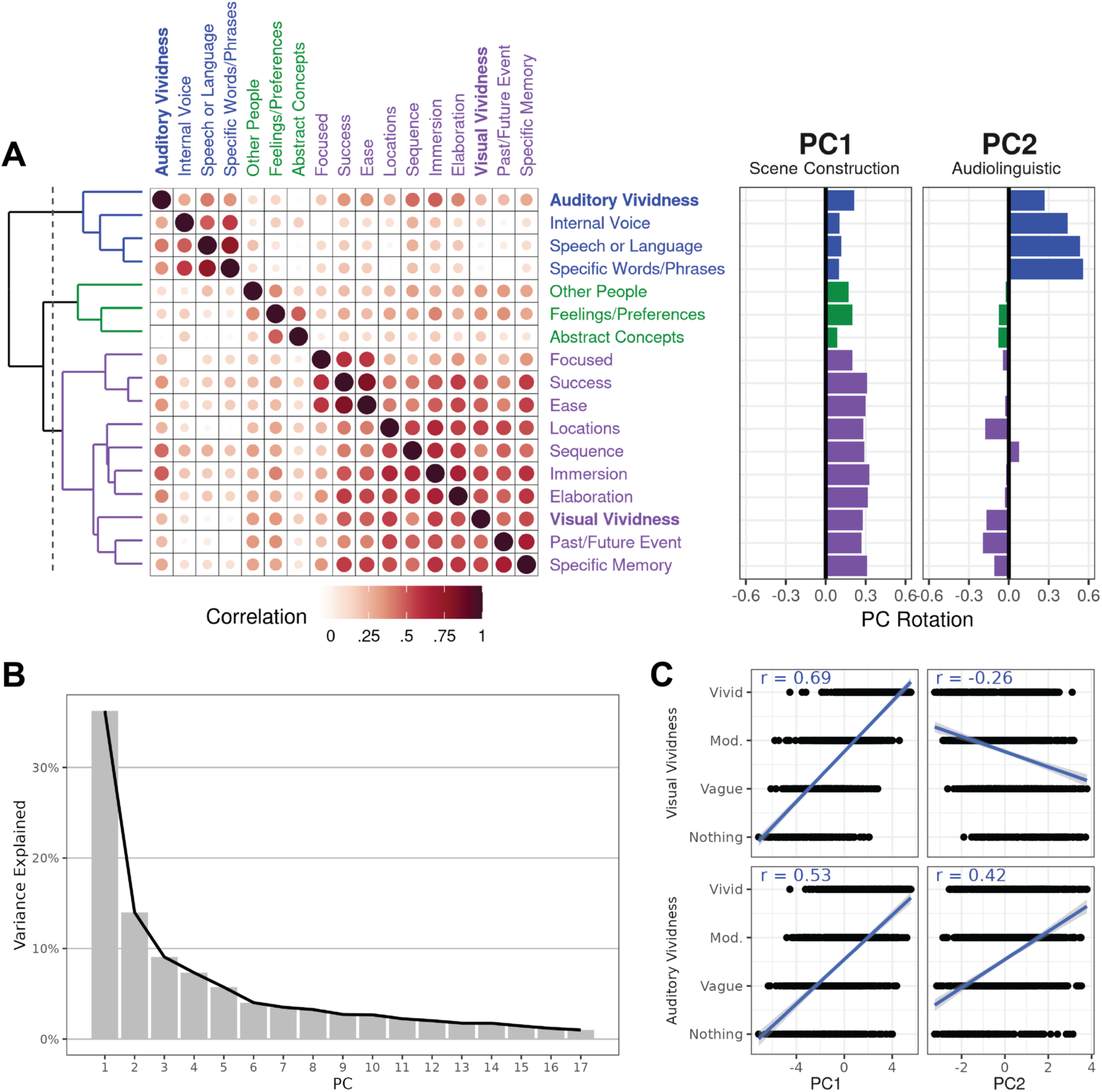
Analysis of follow-up responses reveals multi-dimensional structure of imagined content. A) A correlation matrix was created using all participants’ follow-up responses, organized into clusters using hierarchical clustering (left). We selected the level shown by the dashed vertical line, which defined 3 clusters. The same data was subjected to a principal components analysis (PCA) which revealed two major principal components (right). The rotation of each probe onto the PCs recapitulated the structure seen in the hierarchical clustering analysis. Based on these loadings, we named PC1 “Scene Construction” and PC2 “Audiolinguistic” for convenience. B) A scree plot of the PCA, showing that PC1 and PC2 both explained > 10% of the variance. C) The two components also diJerentiated auditory and visual vividness ratings: PC1 shows a strong positive correlation with both visual and auditory vividness, while PC2 shows a positive correlation with auditory vividness and a negative correlation with visual vividness.

The second component (PC2) explained 14.0% of the variance and differentiated the auditory and language-related probes, including Auditory Vividness, Speech/Language, Inner Speech, and Specific Words/Phrases (see blue bars in Fig. 3A, right), from the scene-construction-related probes (particularly Visual Vividness, Locations, Past/Future Events, and Specific Memory). We therefore refer to PC2 as representing trials high in ‘Audiolinguistic’ processes for brevity, though note it also represents the difference of these trials to the trials rated as high in Scene Construction (i.e., PC1). Accordingly, PC1 showed strong positive correlations with both visual and auditory vividness ratings (Fig. 3C; *r* = 0.69 and 0.53, respectively), while PC2 was negatively correlated with visual vividness and positively correlated with auditory vividness (*r* = -0.26 and 0.42, respectively). Importantly, PC2 did not have high scores for the probes relating to task success (Ease, Success, Focused), so the difference between high and low PC2 loadings likely represents differences in the specific imagined content rather than general task performance. A more fine-grained breakdown of the relationships between PC values and individual thought probes and prompt categories is shown in Supp. Figs. S2 & S3. Importantly, the clustering results emphasized that the task successfully generated mental states that varied in visual and auditory imagery (i.e., PC1 vs. PC2). When participants reported thinking about scenes and events, they typically reported higher visual vividness of their mental states, and when they reported thinking about hearing speech or using inner speech they typically reported higher auditory vividness.

The hierarchical clustering analysis showed a similar pattern to the PCA (see dendrogram in Fig. 3A, left), confirming that the clusters revealed by the PCA were robust across analysis choices, and likely represent true patterns in the responses. Fig. 3A (left) shows the hierarchical clustering level which defined three major clusters, selected as these aligned well with the results of the PCA (i.e., compare the dendrogram in Fig. 3A, left, with the PC loadings in Fig. 3A, right). The first hierarchical cluster resembled the Audiolinguistic PC (i.e., PC2), and comprised the 3 language-related probes plus the auditory vividness probe. A second cluster, which we named “Social”, comprised the probes regarding how much participants thought about Other People, Abstract Concepts, and Feelings/Preferences.

The third cluster resembled the Scene Construction PC (i.e., PC1), and contained the remaining probes including those regarding how much participants thought about Locations, Past/Future Events, a Specific Memory, and probes concerning task success (Immersion, Ease, Success) and other experiential aspects (e.g., how much they elaborated on the prompt, and how much their thoughts were in the form of a sequence of events). This third cluster also included the Visual Vividness responses. Note that in the next level of the dendrogram, which would have defined 4 clusters, the prompts referring to task success (Focused, Success, and Ease) were separated from the scene and event-related probes (e.g., Visual Vividness, Locations, and Past/Future Events).

Thus, the hierarchical clustering confirmed that auditory and visual vividness were separated across task trials and were associated with different types of mental states: auditory vividness was higher when participants thought about speech or language, and visual vividness was higher when participants thought about locations and past or future events.

### Default network A overlaps partially with activity supporting the perception of scenes

The extensive fMRI data collected allowed the estimation of brain networks and task activity patterns within each individual. We defined large-scale networks *a priori* in each individual using functional connectivity (FC) analysis of the independent passive fixation (i.e., “REST”) data. The FC-based networks provided a means to differentiate unimodal and transmodal cortex: unimodal networks are usually constrained to bilateral primary and secondary sensory areas, including the bilateral dorsal and ventral visual streams, whereas the transmodal networks are widely distributed across frontal, parietal, temporal, and midline association areas (e.g., see Yeo et al., 2011; Mesulam, 1998; Margulies et al., 2016; Felleman & Van Essen, 1991; Du et al., 2024).

Surface-projected data were clustered together into 14 or 15 networks using a data-driven approach, and two distributed networks were selected for further analysis, default network A (DN-A; Braga & Buckner, 2017) and the language network (LANG; Braga et al., 2020; Fig. 4) based on findings implicating DN-A in mental scene construction (DiNicola et al., 2023) and LANG in transmodal language processing (Braga et al., 2020; Salvo, Anderson et al., 2024). Based on anatomical landmarks, we also selected networks that covered visual (i.e., visual network, VIS-P, and dorsal attention network, dATN; Corbetta & Shulman, 2002) as well as auditory (AUD) and somatomotor (SMOT) regions.

**Fig. 4:**
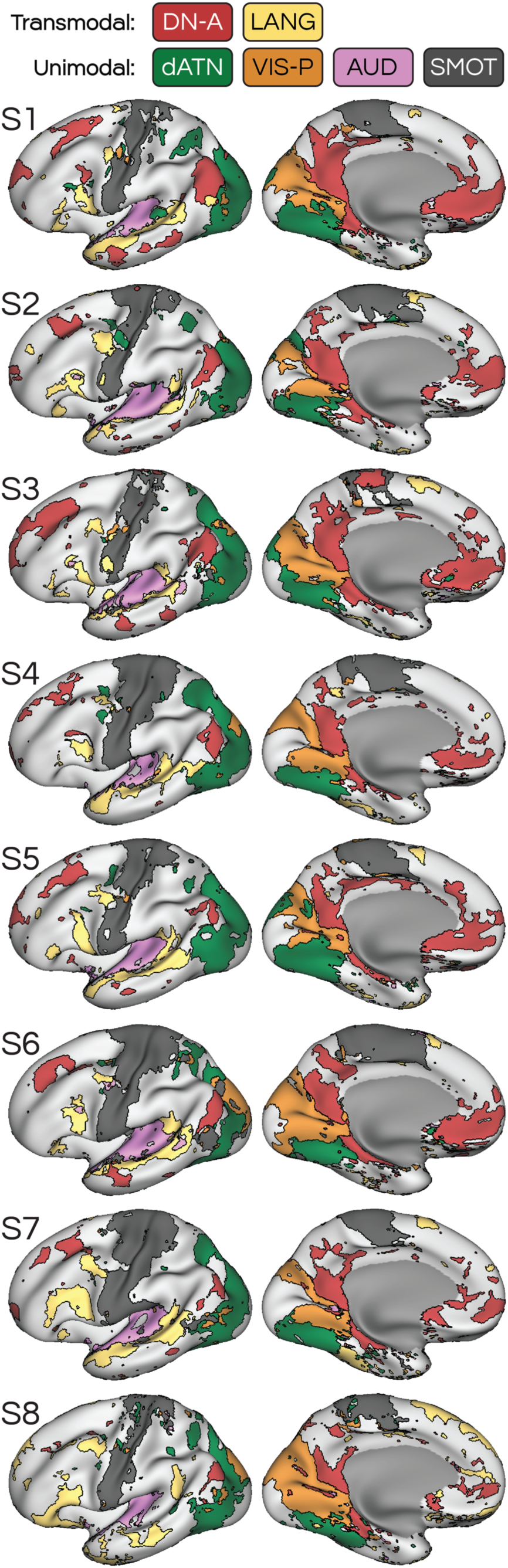
Functional connectivity using independent resting-state data was used to define large-scale brain networks in each participant. Among the 14-15 networks defined for each participant, default network A (DN-A) and the language network (LANG) were identified, based on their anatomical distribution, and selected for further analysis. Additional networks were selected to represent visual, auditory and somatomotor unimodal networks, including dorsal attention network B (dATN), peripheral visual network (VIS-P), primary auditory cortex (AUD), and somatomotor network (SMOT).

In each individual, the FC-defined networks were compared with the task activation maps from the visual perception task, VISCAT, which involved viewing images of different categories (scenes, faces, etc). Supp. Fig. 4 shows that network DN-A overlapped partially with activity evoked by viewing scenes, primarily in parahippocampal and posterior cingulate or retrosplenial cortex. Notably, viewing scenes activated several more posterior regions at or near the occipital lobe that overlapped with the visual processing hierarchy as defined by the VIS-P and dATN networks (Fig. 4; see Felleman & Van Essen, 1991 and Yeo et al., 2011). We explore this later. However, in all cases the scene perception activity extended anteriorly into regions defined as being within DN-A in each participant. Thus, the network maps support that the most anterior regions of the scene-viewing visual stream are actually within transmodal association cortex as estimated by FC.

### The language network overlaps with activity supporting the perception of speech

We recently showed that the LANG network, as defined using FC, encapsulates transmodal language functions in that it can be robustly activated by both auditory and visual forms of language (i.e., listening to speech and reading; Salvo, Anderson et al. 2024). Importantly, the borders of the LANG network along the superior temporal cortex delineated between auditory and transmodal language functions (Salvo, Anderson et al., 2024), reinforcing the division between AUD and LANG evident in this zone (Fig. 4). Thus, similar to the relationship between VIS-P, dATN, and DN-A in regard to scene-viewing, listening to speech similarly activated a set of regions that overlapped unimodal auditory cortex (i.e., AUD) but also extended into the transmodal LANG network (see Supp. Fig. S4). In contrast to the scene-viewing results, which activated part of DN-A only, activity during listening to speech broadly activated the majority of the LANG network.

### Trial-wise experience sampling reveals more widespread brain activity compared to condition-wise analysis approach

We next computed the vertex-wise responses during each trial of the IMAGINE task for each individual to investigate brain activity during self-generated states. Three analysis approaches were taken (Supp. Fig. S5). First, we grouped trials according to the original categories designed into the experiment, including Scenes, Faces, Sounds, Perceptual Speech, and Inner Speech conditions. Trial-related responses were averaged together within each category, and then contrasted across categories. The Sounds condition was chosen as the contrast condition to control for basic task demands (e.g., reading, attending to the screen), while serving as a common baseline for the Scene and Language-related conditions. The Scenes > Sounds contrast maps revealed that focal regions at or near the parahippocampal cortex consistently showed increased activity during imagining scenes for nearly all subjects except S6 (Supp. Fig. S6, left). Beyond this parahippocampal region, some subjects also showed evidence of activity at or near the retrosplenial cortex, and a few (e.g., S5, S8) showed activity in other regions including the posterior inferior parietal lobe.

The maps of imagining-related activity calculated using the language conditions (contrast of Perceptual Speech and Inner Speech > Sounds) revealed task-evoked responses across the lateral surface (Supp. Fig. S6, right). However, these condition-wise maps were inconsistent, with many subjects showing few active regions (e.g., S7, S8), while others showed clear activity overlapping the LANG network (e.g., S3, S6). This heterogeneity might be because participants varied across trials in how they imagined each prompted content, even when prompts were from the same condition. Analysis of the experience sampling data showed that in many instances, a prompt that was intended to elicit one type of mental state often elicited others (e.g., see Fig. 1 and Supp. Fig. S1). For example, in Supp. Fig. S7, it is shown that participants rated the Perceptual Speech condition as being higher in visual vividness, presumably because many of the prompts (e.g., “Imagine a couple arguing”) elicited integrated audiovisual mental content. Conversely, participants generally reported the Inner Speech condition as being higher in terms of whether they thought about specific, identifiable words. These differences underscore the limitations of a condition-wise approach, and provide strong motivation to explore brain activity using trial-wise analyses that can account for such variation.

We undertook two strategies to account for trial-wise phenomenology as captured by the mDES surveys (Fig. 3): an *a priori* selected composite score and a data-driven clustering approach. First, we computed two composite scores, following DiNicola et al. (2023), targeting the clusters shown in Fig. 3A. We contrasted trials scoring in the top 20% with the lowest 20% of each composite. The first composite score, consisting of a sum of the Visual Vividness and Locations thought-probe ratings, revealed more robust activity, which recapitulated the condition-wise activity within parahippocampal regions and also included more extensive activity within the retrosplenial and posterior cingulate cortex and other regions of DN-A (including in the inferior parietal lobe and rostral medial prefrontal cortex; Supp. Figs. S5 & S8).

The maps for the language composite scores, which consisted of a sum of the Auditory Vividness, Speech/Language, and Specific Words/Phrases thought-probes, showed an improvement in some subjects (e.g., S2, S3, possibly S6), but other subjects showed similar activity patterns to the condition-wise analysis (Supp. Fig. S5, lower middle column; Supp. Fig. S8, right column).

Finally, we accounted for the trial-wise variance in phenomenology more comprehensively by incorporating the principal component loadings from the PCA (Fig. 3) that considered all the mDES responses. The trial-wise loadings from PC1 and PC2 were mean-centered and entered into a general linear model, and at each vertex a z score was calculated representing how much that vertex’s activity in each trial changed as a function of how the loadings for PC1 and PC2 changed across trials (i.e., the relationship between PC loadings and trial-evoked activity). These maps provided an improvement on the condition-wise and composite score analyses (Supp. Fig. S5), and were used for the remaining analyses.

Notably, only the condition-wise analyses revealed activity for scene imagery that was restricted to the parahippocampal cortex. In contrast, accounting for phenomenological reports revealed much more distributed activity throughout the cortex (Fig. 5), suggesting network-level activation during the imagining of scenes (DiNicola et al., 2023). These comparisons strongly support that conventional condition-wise analyses fail to capture important aspects of participants’ mental states that can be retrieved using experience sampling and leveraged to better reveal brain activity related to those states.

**Fig. 5:**
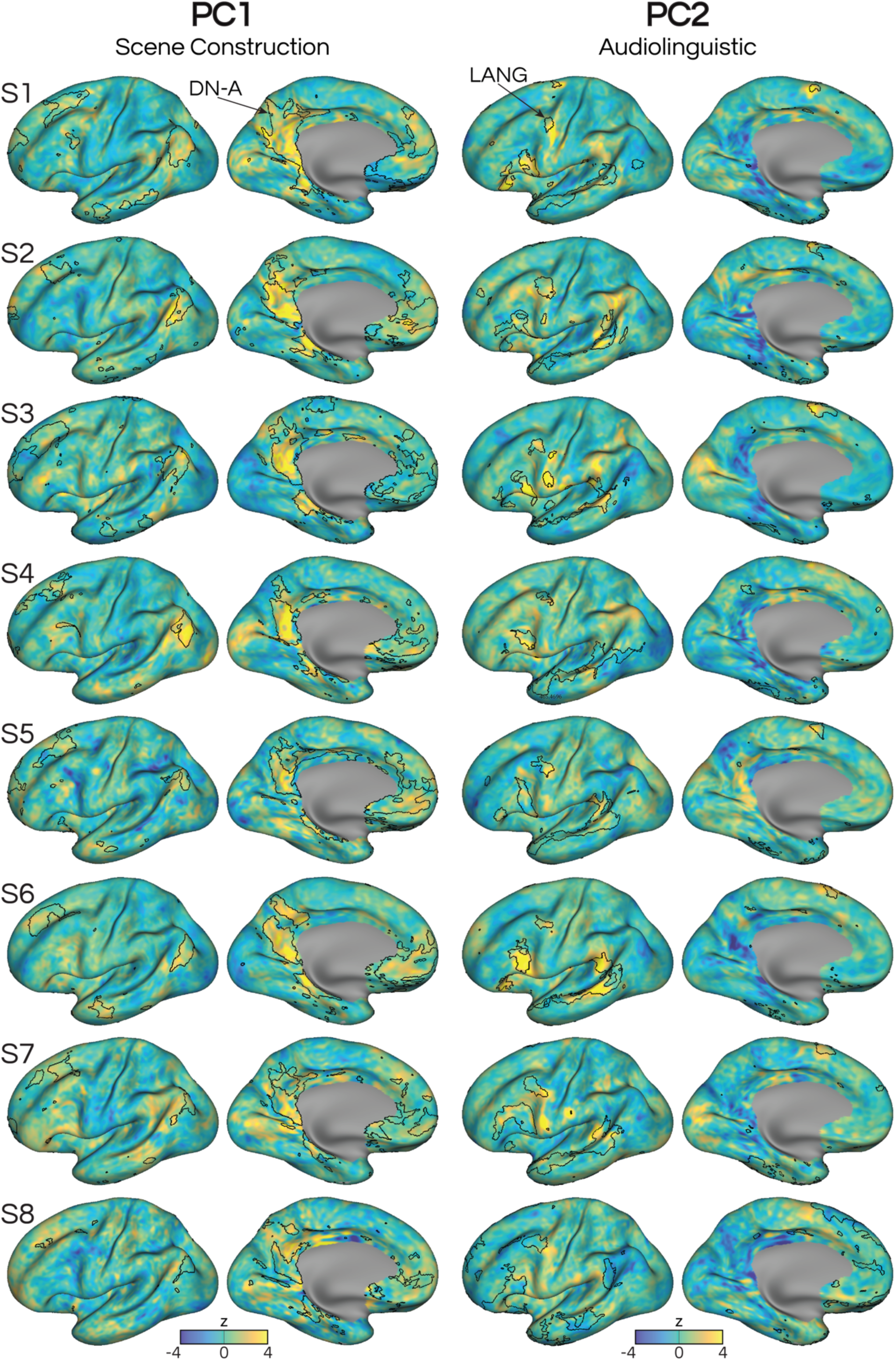
Accounting for trial-wise experience sampling responses reveals widespread activity within DN-A and LANG during imagining scenes and speech, respectively. At each vertex, a GLM was created using trial-wise loadings for both the Scene Construction (PC1) and Audiolinguistic (PC2) factors (see Fig. 3). The resulting map for PC1 shows overlap primarily with DN-A, including in anterior regions such as the medial prefrontal and dorsolateral prefrontal cortex. The map for PC2 shows positive regions that overlap with LANG, though the overlap was clearer in some subjects (e.g., S3, S5, S6) than others. The PC2 map showed negative regions that likely overlapped with DN-A, which were in line with PC2’s negative scores for the Locations and Visual Vividness related mDES survey responses (Fig. 3).

### Different imagined states recruit different distributed brain networks

Analysis of brain activity related to the two principal components (PC1 & PC2) revealed that mental imagery was linked to activity in two separate networks, DN-A and LANG (Fig. 5). For PC1, scene-imagining-related activation was located within the boundaries of DN-A, including parahippocampal cortex, retrosplenial and posterior cingulate cortex, inferior parietal cortex, and rostromedial prefrontal cortices (Fig. 5, left column). In contrast, for PC2, which represented the Audiolinguistic factor, activation was observed in multiple parts of the LANG network, including lateral temporal cortex, inferior and middle frontal cortex, and the supplementary motor area – especially in subjects S1, S2, S3, S6, and S7 (Fig. 5, right column). The PC2 maps were not as robust in the remaining subjects, showing generally lower magnitude and patchier regions of activation.

The Audiolinguistic factor may have led to less robust maps than the Scene Construction factor as a result of more variable activity during relevant task conditions. The direct contrast of the Perceptual Speech and Inner Speech imagination conditions (Supp. Fig. S7) revealed some systematic differences in patterns of brain activity: Inner Speech showed greater activity at or near posterior superior temporal cortex and precentral cortex, which were close to the LANG network and ventral motor strip, which putatively could be at or near regions controlling articulation (de Heer et al., 2017; Krubitzer, 2007). Meanwhile, Perceptual Speech generally showed greater activity in posterior parietal regions such as posteromedial (i.e., retrosplenial and posterior cingulate) and parahippocampal cortices that appeared to more closely resemble DN-A and activity patterns related to imagining scenes (Fig. 5). These differences between the language-related prompt conditions argue that different strategies for imagining speech, such as imagining hearing someone else speak versus thinking of specific words in inner speech, may have been implemented by participants across different trials (Sulfaro et al., 2024; Alderson-Day & Fernyhough, 2015; Carruthers, 2018). This might have led to higher variability in activity related to the Audiolinguistic PC2 compared to the Scene Construction PC1 (Fig. 5).

Notably, activation related to PC2, which distinguished the Audiolinguistic probes with positive scores and Scene Construction probes with negative scores (see Fig. 3A, right), also revealed negative z scores in DN-A regions, including the parahippocampal cortex and retrosplenial cortex, in many participants (Fig. 5). Thus, PC2 also demonstrates that mental states described as being high in Audiolinguistic versus Scene Construction content activate distinct distributed networks, suggesting that recruitment of DN-A during trials high in Scene Construction loadings is not simply due to a general task performance effect of the imagining task (i.e., PC2 did not have strong loadings for the performance- and success-related thought probes).

We quantified these results by assessing the distribution of z scores for all vertices within the two *a priori* defined transmodal networks, DN-A and LANG (Fig. 4). In all subjects (even those that showed few positive regions in the whole-brain maps in Fig. 5), z scores for each PC were dissociated between the two networks: DN-A showed higher responses during Scene Construction-related trials, while LANG showed higher responses during Audiolinguistic-related trials. Across all subjects, PC1 values within DN-A were consistently higher than those within LANG (two-tailed t-test, all *p*s < .001 after Bonferroni correction; Fig. 6, left column). For PC2, values within LANG were consistently higher than those within DN-A in each subject (two-tailed t-test, all *p*s < .001 after Bonferroni correction; Fig. 6, right column). Thus, the results support that distinct distributed networks are recruited during these different mental states.

**Fig. 6:**
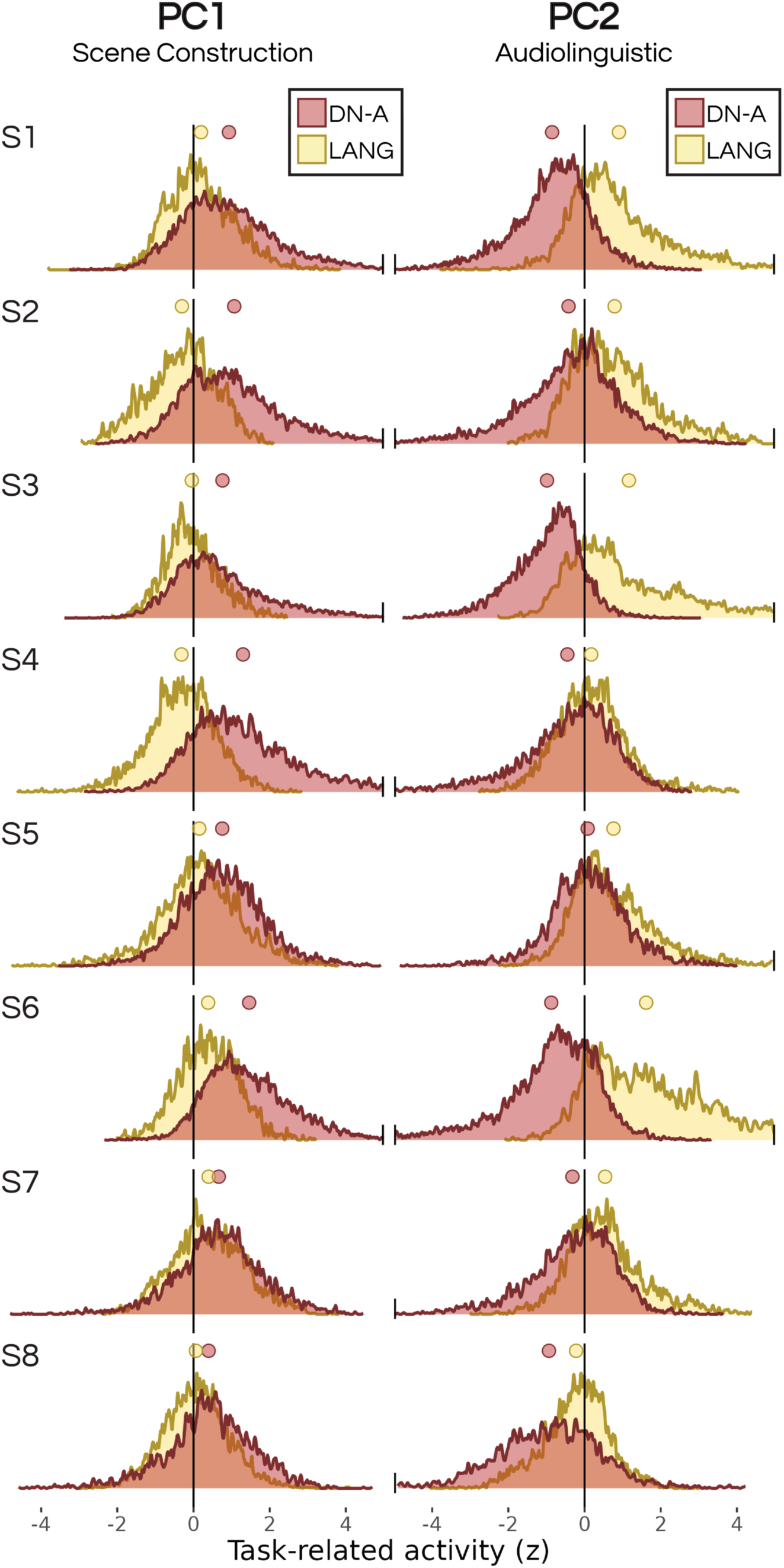
Imagining scenes versus language reinstates activity in diTerent large-scale association networks. The distribution of z-scored beta values from within each transmodal network, DN-A and LANG, are shown, calculated from the regression of PC1 and PC2 against brain activity (see full maps in Fig. 5). Values are plotted using normalized probability density functions. The average values for each network are represented with a colored circle above the distribution. In all cases, the distribution of z values for DN-A is higher than LANG for PC1, while the inverse is true for PC2. Values are truncated at +/- 5 for visualization purposes.

### Imagination and perception overlap within transmodal association networks

A final set of analyses leveraged the network mapping data to determine whether regions that were recruited by both imagined and perceptual states were located within unimodal or transmodal networks. We thresholded and binarized the task maps of viewing scenes (Supp. Fig. S4) and imagining scenes (Fig. 5) and computed their overlap map. Along the ventral temporal cortex, the more anterior parts of the scene-perception stream overlapped with activity during imagining scenes. These overlap regions were at a location further away from unimodal visual areas, and further into association cortex, and were located within DN-A as mapped within most individuals (Fig. 7). This is interesting, as the canonical DN is not conventionally thought to be involved in perception, and indeed shows reduced activity during externally oriented perceptual tasks (Spreng et al., 2010; Shulman et al., 1997). Here, we show that when networks are mapped within individuals, the borders of DN-A encompass the regions involved in imagining scenes, and that regions active during viewing scenes extend only partially into DN-A. This agrees with previous work comparing perception and mnemonic representation of scenes (Silson et al., 2019b; Steel et al., 2021, 2023).

**Fig. 7:**
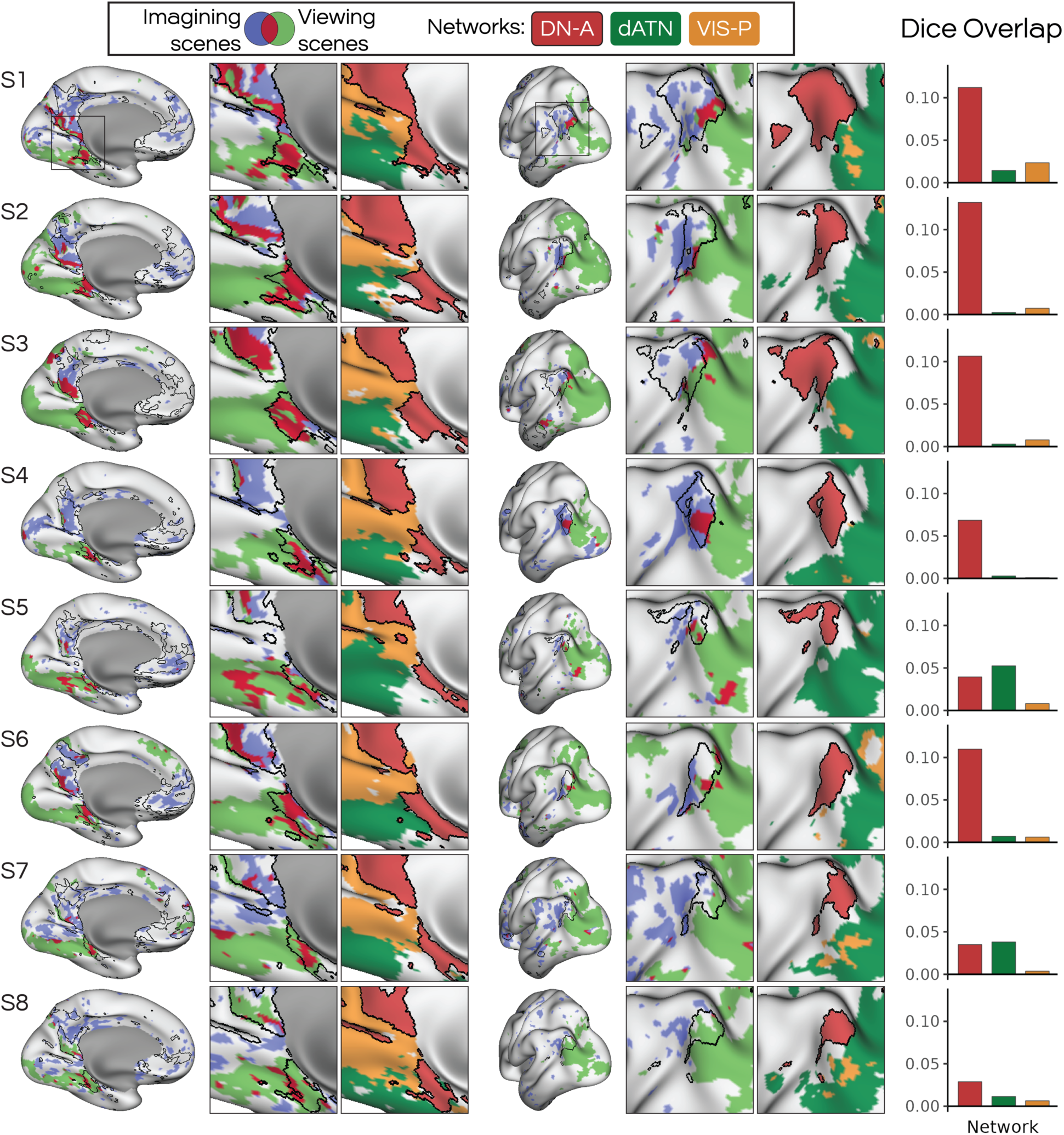
Imagining and viewing scenes overlapped within the boundaries of transmodal default network (DN-A). Surface-projected maps show regions active during imagining scenes in blue (from Fig. 5, thresholded and binarized) and regions active while viewing scenes in green (from Supp. Fig. S4; thresholded and binarized). Regions that overlapped between the two are shown in red. The insets show a zoom-in on regions of overlap, showing the task overlap map alongside a zoom in of the functional networks defined in each individual (see full networks in Fig. 4). Imagining scenes shows overlap with scene viewing in parahippocampal and retrosplenial cortices, and this overlap was predominantly within the bounds of network DN-A, not adjacent visual networks, as shown by the dice coeJicients plotted on the right. Other networks shown are the dorsal attention network (dATN), and peripheral visual network (VIS-P).

Additionally, we noted that the overlap between perception and imagining within DN-A was seen largely near the borders between DN-A and adjacent networks involved in visual perception, the dorsal attention network and peripheral visual network (shown in Fig. 4; see Du et al. 2024). These findings suggest that the confluence between these networks may be an important factor in understanding why these regions are recruited during imagining of scenes (Steel et al., 2025). But importantly, the overlap between imagining and perception of scenes was often confined to the regions of DN-A (see especially subject S3 in Fig. 7).

Our calculation of Dice coefficients demonstrates that, for 6 out of 8 subjects, the regions overlapping imagery and perception were predominantly within DN-A compared to these adjacent, visual networks.

A similar observation was made for the overlap between imagining and listening to speech (Fig. 8). Despite this type of imagining being different in phenomenological content (Fig. 3), and despite the focus being on a different large-scale network, again the overlap between imagining and perceiving was found to be largely confined to the bounds of the transmodal network, LANG (e.g., see especially S6 in Fig. 8). Listening to speech robustly activated the LANG network, as well as nearby auditory areas. However, of these regions, those that fell within the LANG network were more likely to be reinstated when participants imagined speech. Similar to imagining scenes, this overlap tended to occur near the boundaries between LANG and adjacent sensory areas encapsulated by the auditory (AUD) and somatomotor (SMOT) networks (Fig. 8). Activity related to imagining speech was also located farther away from unimodal (auditory or motor) cortices, extending further into association cortex. The dice coefficients showed that in all 8 subjects the overlap between imagining and listening to speech was predominantly within LANG regions rather than AUD or SMOT networks. Thus, the speech imagery results reiterated the findings from the scene imagery trials, emphasizing that the self-generated states elicited here recruited transmodal distributed networks, rather than the adjacent sensory areas.

**Fig. 8:**
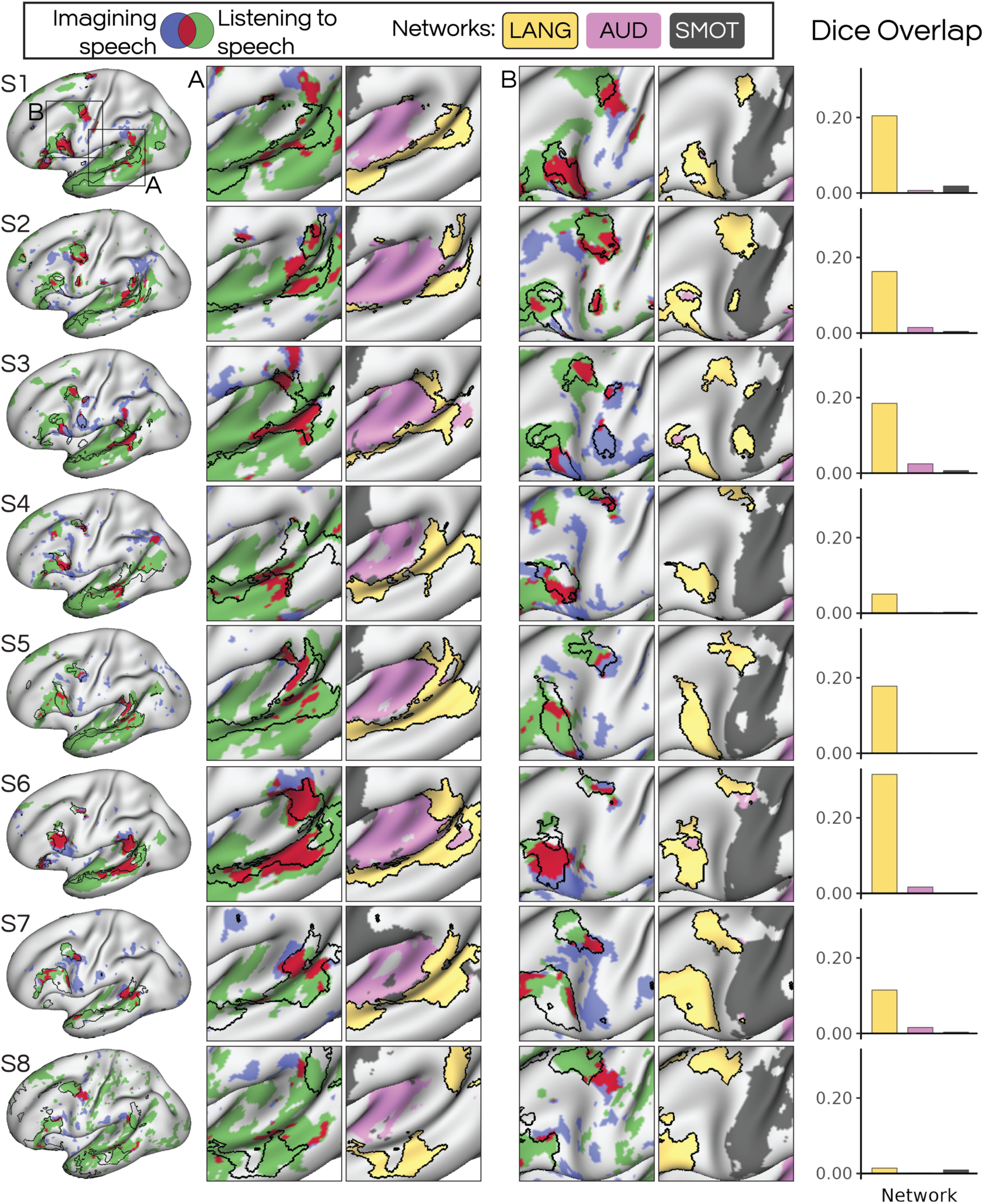
Imagining and listening to speech overlapped within the boundaries of the transmodal language network (LANG). Surface-projected maps show regions active during imagining speech in blue (from Fig. 5, thresholded and binarized) and regions active while listening to speech in green (from Supp. Fig. S4; thresholded and binarized). Regions that overlapped between the two are shown in red. The insets show a zoom-in on regions of overlap, alongside a zoom in of the functional networks defined in each individual (see full networks in Fig. 4). Imagining speech shows overlap with listening to speech along the length of lateral temporal cortex, in regions that encircled the auditory (AUD) network. Overlap was also seen in the prefrontal cortex, including inferior and middle frontal cortex, that again was positioned at the interface between LANG and the somatomotor network (SMOT). Regions of overlap fell predominantly within the distributed LANG network, not adjacent AUD and SMOT networks, as shown by the dice coeJicients plotted on the right.

## Discussion

This study showed that imagining of different types of content is associated with the activation of different large-scale networks and explored the extent to which brain regions involved in perception are reactivated to support imagined states.

Using a novel imagination task, we were able to elicit self-generated mental states that varied in their vividness of visual and auditory imagery (Fig. 1) and content (Fig. 3). By collecting extensive self-report data from each participant using a multi-dimensional experience sampling approach (mDES), we showed that trials in which participants reported thinking about locations and/or events were associated with high vividness of visual imagery (Fig. 3; see also DiNicola et al. 2023). In contrast, trials in which participants reported thinking using speech and language were associated with high vividness of auditory imagery (Fig. 3). By leveraging a precision fMRI approach, where functional anatomy is defined within individuals (Fig. 4), we showed that these different types of mental states were linked to the activation of different large-scale networks, DN-A and LANG (Fig. 5 & 6). Notably, the use of self-report mDES data was critical to resolving mental state-related activity, and improved on a conventional ‘condition-wise’ approach that assumed that participants engaged with each imagination prompt as intended in each condition (Supp. Fig. S3). When we accounted for self-reported mental content in each trial, we observed that imagining scenes and speech were associated with widespread activity within multiple regions of DN-A and LANG, respectively (Fig. 5). These findings support network-level engagement during self-generated mental states.

### Transmodal distributed networks are active during imagining

A core distinction in brain organization is the separation between networks serving unimodal functions (e.g., processing sensory information), and networks serving associative or transmodal functions (Mesulam, 1998). Language is an intuitive example of a transmodal function, in that the same concepts can be conveyed via visual (e.g., written) or auditory (e.g., spoken) words. In each case, shared cognitive processes are likely to underlie the translation of sensory stimuli into their associated concepts according to syntactic rules (Fedorenko et al., 2024). Previous work has shown that the LANG network, as defined using FC, likely encompasses such transmodal functions, as it shows robust responses during both reading and listening to speech (Salvo, Anderson et al., 2024; see also Scott et al., 2017; Fedorenko et al., 2010; Yuan et al., 2023). Similarly, DN-A can be activated by tasks where participants disengage from the external world (Smallwood et al., 2009; Shulman et al., 1997; Raichle et al., 2001), but also tasks targeting processes such as navigation and the consideration of spatial relationships (Peer et al., 2015; DiNicola et al., 2023; Silson et al., 2019b). These findings support a transmodal role of DN-A that is not strictly tied to visuospatial (i.e., visual scene-related) processing. Furthermore, the organization of both LANG and DN-A, where each network contains regions in multiple cortical association zones, fits the distributed network motif that is characteristic of association cortex (Goldman-Rakic, 1988; Mesulam, 1990; 1998), and is fundamentally different from the hierarchically organized regions that serve sensory processing (Felleman & Van Essen, 1991). Thus, LANG and DN-A both likely represent transmodal associative, rather than sensory-specific, functions.

A main finding here is that imagining speech or scenes mainly recruited the transmodal association networks, rather than the adjacent unimodal sensory networks (Fig. 5). This finding was consistent for thinking about scenes and speech, though in each case different distributed networks (DN-A and LANG) were involved. Further, activity in the two networks was directly related to participants’ trial-wise assessments of the visual and auditory vividness of their imagined content (Fig. 3A & 3C). Prior work has suggested that brain activity related to imagined states follows the principle of *sensory reinstatement*: the recruitment of sensory regions in the absence of sensory input. In our study, the within-individual network maps supported that self-generated states are more likely to elicit activity within the distributed networks (DN-A, LANG) rather than in unimodal sensory (VIS-P, dATN, AUD) networks. The boundaries of the distributed networks often encapsulated the regions active during imagining (Fig. 5), and the overlap between imagining and perception was predominantly within the distributed networks, not adjacent sensory networks (Figs. 7 & 8). This is a departure from the sensory reinstatement account: our work supports it is the transmodal regions that are “reinstated” during imagined states (Doucet et al., 2012). Conversely, it is interesting to reflect that perceptual processing may in fact involve a combination of unimodal and transmodal areas, the latter of which are reinstated for imagined states.

### Perception and imagination overlap at the interface between unimodal and transmodal networks

Our results suggest that the perception of scenes and speech involves multiple regions that include sensory (visual, auditory) areas but also extend into association cortex (Supp. Fig. S4; Figs. 7 & 8), and therefore cut across the unimodal-transmodal distinction. Perceiving scenes and speech evoked responses within visual and auditory cortex, respectively. Scene-viewing-related activity extended from the occipital pole and radiated along the ventral temporal cortex in a posterior-anterior arc, culminating in what is likely to be the parahippocampal place area (Supp. Fig. S4; Kanwisher, 2010). This more anterior region overlapped with DN-A in all subjects (Supp. Fig. S4; see also Steel et al., 2025; Silson et al., 2019a). Speech-listening-related activity extended from auditory areas at or near Heschl’s gyrus, in the upper bank of the superior temporal sulcus within the sylvian fissure, and radiated outwards to surrounding lateral temporal cortex regions. These additional regions overlapped with the LANG network in all subjects (Supp. Fig. S4). Thus, both perceptual tasks led to activation of unimodal sensory networks (i.e., VIS-P, dATN, AUD), but also extended to transmodal networks (i.e., DN-A, LANG).

We recently showed that, in these same subjects, the boundaries of the LANG network along the superior temporal cortex, as mapped using intrinsic functional connectivity (Fig. 4), provide a relatively accurate proxy for the separation between auditory functions (i.e., responding to sounds not containing comprehensible language) and transmodal language functions (i.e., regions responding to both visual and auditory sentences; Salvo, Anderson et al., 2024). Thus, task-based analysis in these same participants supports that the side-by-side regions of the intrinsic FC-defined AUD and LANG networks represent the separation between unimodal and transmodal functions. Thus, listening to speech appears to activate both unimodal and transmodal regions as one moves away from auditory cortex. Similar findings have been observed in the scene-viewing literature: the more anterior regions of inferior parietal, posteromedial and ventral temporal cortex have more mnemonic roles compared to their adjacent, more posterior, more visual counterparts. The more posterior regions are more strongly activated by viewing scenes, while the more anterior parts are more activated by remembering familiar places (Steel et al., 2021). Here, we show that this distinction between imagined and perceptual processing of scenes seems to accord with the boundaries dividing DN-A and vision-related networks dATN and VIS-P (Fig. 7; Steel et al., 2025). Thus, the interface between transmodal and unimodal networks appears to hold key value in understanding the relationship between perception and imagination, both in the visuospatial and audiolinguistic domain.

While we observed activation in early visual regions during perception of scenes, and in early auditory regions for language perception, we did not see this for imagining. This omission could be due to the specific instructions given to participants, which directed them to imagine using broad cues (e.g., imagine a castle on a hill), without specific instructions as to the lower-level details (e.g., orientation, visual field location, color, tone, etc.) of the imagined content. Such broad imagining has been shown to solicit high ratings of subjective vividness, even without lower-level imagined features (Bigelow et al., 2023). One proposal is that self-generated states that are focused on lower-level features may more readily reveal activity within unimodal areas that actually encode those features (Dijkstra, 2024). Thus, our results do not negate the idea that reinstatement of earlier sensory regions could be a correlate of mental imagery: it is possible that mental imagery involves the activation of much of the visual or auditory sensory hierarchies, and including downstream transmodal areas, but earlier stages of those hierarchies were not revealed by our analyses because lower-level sensory features were not consistent across trials. Nonetheless, our results suggest that commonly reported self-generated states involving wholistic, integrated content, such as imagining scenes and speech, are inextricably linked to the activation of large-scale networks (Fig. 5). Further, our results support that activation of the large-scale networks is directly correlated with participants’ self-reported vividness of their imagined content, such that stronger activity in LANG was linked to higher auditory vividness, and stronger activity in DN-A was linked to higher visual vividness. Thus, our results refine the idea that the key correlate of mental imagery is reinstatement of sensory regions: here the clearest activation was found in transmodal association networks, and the regions that overlapped between imagery and perception were largely confined to those transmodal networks (Figs. 7 & 8).

### Variation across subjects suggests heterogeneity in correlates of inner speech

Our maps of activity related to imagining speech were more variable across subjects than for imagining scenes. Speech comprehension requires holding words in working memory and combining these with incoming words which can alter word- and phrase-level meaning (Lewis et al., 2006). One proposal is that the LANG network represents ‘core’ language functions required for linking combinations of sensory percepts (e.g., heard or read words) with their associated meanings according to syntactic rules (Fedorenko et al., 2024). Imagining hearing someone speak may not elicit the same demands on these combinatorial/syntactic processes compared to hearing actual speech. For instance, a participant may focus on the tonal qualities of an imagined voice, rather than imagining specific, identifiable words being uttered. It could be that the extent to which participants activated the LANG network during the imagination task reflects the extent to which they imagined real words and sentences. Neither the Perceptual Speech condition (e.g., “Imagine hearing a teacher talk”) nor most prompts from the Inner Speech condition (e.g., “Imagine listing the names of fruits”) demanded imagining syntactic combinations of words, and so may not have targeted the core functions of the language network (the full list of prompts is provided in the supplementary materials). When we studied the mDES responses we saw that, while all participants reported greater specific word content for the two speech categories overall, almost all participants showed multiple speech trials in which they did not endorse imagining specific words or phrases (Supp. Fig. S7).

Prior work has suggested that spontaneous speech imagery is more likely to lead to activation of auditory areas, compared to the type of cued imagining used here (Hurlburt et al., 2016). Some prior work has shown overlap between perceiving and imagining speech in classic language areas such as the lateral temporal cortex, inferior frontal gyrus, middle frontal gyrus, and the supplementary motor area (Lu et al., 2023; Stephane et al., 2021; Yao et al., 2011; Perrone-Bertolotti, 2012; see also Kleber et al., 2007). However, several studies have suggested that different forms of imagining speech (e.g., imagining articulating versus hearing speech) are supported by different brain regions (Tian et al., 2016; see also Pratts et al., 2023; Paulesu et al., 1993). Along this theme, Jackendoff (1996) proposes that inner speech is primarily auditory and does not involve concepts: the meaning of spoken words is a separate “conceptual structure” that is unconscious, whereas inner speech is auditory and conscious. In this framework, inner speech is the phonological structure linked to the concept (Jackendoff, 2011), and would hence be related to auditory imagery. Others propose that inner speech involves a sub-articulatory preparatory motor command that is not acted upon (Fernyhough and Borghi, 2023; Pratts et al., 2023). While we did not design the present study to address this, our comparison of Perceptual Speech and Inner Speech conditions showed that the Inner Speech condition, where participants reported a higher incidence of thinking about specific words, in some subjects revealed stronger activity in regions close to the ventral precentral gyrus at or near regions that control the orofacial musculature (Supp. Fig. S7; Krubitzer, 2007). The distribution of regions resembled those revealed by Papoutsi et al. (2009) as being involved in phonetic processing of meaningless but pronounceable pseudowords, and have previously been implicated in articulatory and/or phonemic processing (e.g., Krubitzer, 2007; Hickok & Poeppel, 2007; de Heer et al., 2017). In contrast, in the Perceptual Speech condition, in which participants tended to report thinking more about Locations and experiencing higher visual vividness, we observed increased activity in a set of regions that resembled DN-A (Supp. Fig. S7). This indicated that imagining hearing someone speak was more often associated with visual and scene-related mental imagery than using inner speech, and, perhaps accordingly, was also associated with increased engagement of DN-A.

These factors highlight that variability in the possible forms of imagining speech may be a factor in the variability we observed here in brain activity. In addition, some of our prompts targeted inner speech by asking participants to recite names (e.g., “Imagine the names of US presidents”) or numbers (e.g., “imagine counting up from 5”). These may have led to different patterns of brain activity, explaining the greater heterogeneity seen for the imagined speech trials in Fig. 5. We offer the maps in Supp. Fig. S7 as a hypothesis generating analysis, and note that the maps varied considerably across individuals despite the aforementioned putative consistencies. Additional research, using different sets of mDES survey questions, may be able to disentangle correlates of these putatively different forms of inner speech.

### Limitations and technical considerations

A limitation of the present study is that we were only able to elicit some types of mental states and phenomenological reports, given the constraints of time in and out of the scanner. Additional studies could leverage the task used here to elicit a wider variety of mental states, and to use additional mDES thought probes to investigate other phenomenological features. Another limitation that our results highlighted is that the prompts often evoked integrated mental states that were high in both visual and auditory content (Fig. 1). Some prompts targeting the Sounds condition were able to elicit high auditory vividness with low visual vividness (e.g., “Imagine a rock song playing on the radio”), however the majority elicited multimodal imagery. This may reflect that imagined states are highly integrated, especially when prompted by broad descriptions as used in the present study. Future studies could devise prompts that better separate auditory imagery, such as by targeting music.

In conclusion, we show that self-generated mental states are closely related to network-level activation of large-scale distributed networks that populate association cortex. We further show that activity in these networks is related to the self-reported vividness of visual and auditory imagery. Our results support that brain regions that overlap between imagination and perception of the same types of content actually fall within transmodal networks, not unimodal sensory regions, challenging sensory reinstatement as a key correlate of self-generated mental states.

## Acknowledgements

This research was supported in part through the computational resources and staN contributions provided for the Quest high performance computing facility at Northwestern University which is jointly supported by the Office of the Provost, the Office for Research, and Northwestern University Information Technology.

## Author Contributions

Conceptualization: NLA, JS, RMB

Data curation: NLA, JJS, RMB

Formal analysis: NLA, JJS

Funding acquisition: NLA, JJS, RMB

Investigation: NLA, JJS, RMB

Methodology: NLA, JJS, RMB

Resources: RMB

Supervision: RMB

Visualization: NLA, RMB

Writing – original draft: NLA, RMB

Writing – review & editing: NLA, JJS, JS, RMB

## Data and materials availability

All data needed to evaluate the conclusions in the paper are present in the paper and/or the Supplementary Materials or will be made available upon publication.

## References

B. Alderson-Day, C. Fernyhough, Inner speech: Development, cognitive functions, phenomenology, and neurobiology. Psychol. Bull. 141, 931 (2015).

A. Aleman, E. Formisano, H. Koppenhagan, P. Hagoot, E. H. De Haan, R. S. Kahn, The functional neuroanatomy of metrical stress evaluation of perceived and imagined spoken words. Cereb. Cortex 15, 221–228 (2005).

E. Amit, C. Hoeflin, N. Hamzah, E. Fedorenko, An asymmetrical relationship between verbal and visual thinking: Converging evidence from behavior and fMRI. Neuroimage 152, 619–627 (2017).

J. R. Andrews-Hanna, J. S. Reidler, J. Sepulcre, R. Poulin, R. L. Buckner, Functional-anatomic fractionation of the brain’s default network. Neuron 65, 550–562 (2010).

W. A. Bainbridge, E. H. Hall, C. I. Baker, Distinct representational structure and localization for visual encoding and recall during visual imagery. Cereb. Cortex 31, 1898–1913 (2021).

G. H. Betts, The distribution and functions of mental imagery (Columbia Univ., New York, 1909).

E. J. Bigelow, J. P. McCoy, T. D. Ullman, Non-commitment in mental imagery. Cognition 238, 105498 (2023).

R. M. Braga, R. L. Buckner, Parallel interdigitated distributed networks within the individual estimated by intrinsic functional connectivity. Neuron 95, 457–471 (2017).

R. M. Braga, K. R. Van Dijk, J. R. Polimeni, M. C. Eldaief, R. L. Buckner, Parallel distributed networks resolved at high resolution reveal close juxtaposition of distinct regions. J. Neurophysiol. 121, 1513–1534 (2019).

R. M. Braga, L. M. DiNicola, H. C. Becker, R. L. Buckner, Situating the left-lateralized language network in the broader organization of multiple specialized large-scale distributed networks. J. Neurophysiol. 124, 1415–1448 (2020).

R. L. Buckner, D. C. Carroll, Self-projection and the brain. Trends Cogn. Sci. 11, 49–57 (2007).

R. L. Buckner, J. R. Andrews-Hanna, D. L. Schacter, The brain’s default network: anatomy, function, and relevance to disease. Ann. N.Y. Acad. Sci. 1124, 1–38 (2008).

N. Bunzeck, T. Wuestenberg, K. Lutz, H. J. Heinze, L. Jancke, Scanning silence: mental imagery of complex sounds. Neuroimage 26, 1119–1127 (2005).

G. Cabbai, C. Racey, J. Simner, C. Dance, J. Ward, S. Forster, Sensory representations in primary visual cortex are not sufficient for subjective imagery. bioRxiv 2024–01 (2024).

J. Carroll, Imagination, the brain’s default mode network, and imaginative verbal artifacts. Evol. Perspect. Imagin. Cult. 31–52 (2020).

P. Carruthers, “The causes and contents of inner speech” in Inner Speech: New Voices, P. Langland-Hassan, A. Vicente, Eds. (Oxford Univ. Press, Oxford, UK, 2018).

W. Chen, T. Kato, X. H. Zhu, S. Ogawa, D. W. Tank, K. Ugurbil, Human primary visual cortex and lateral geniculate nucleus activation during visual imagery. Neuroreport 9, 3669–3674 (1998).

R. M. Cichy, J. Heinzle, J. D. Haynes, Imagery and perception share cortical representations of content and location. Cereb. Cortex 22, 372–380 (2012).

M. Corbetta, G. L. Shulman, Control of goal-directed and stimulus-driven attention in the brain. Nat. Rev. Neurosci. 3, 201–215 (2002).

R. W. Cox, AFNI: software for analysis and visualization of functional magnetic resonance neuroimages. Comput. Biol. Med. 29, 162–173 (1996).

R. W. Cox, AFNI: what a long strange trip it’s been. Neuroimage 62, 743–747 (2012).

X. Cui, C. B. Jeter, D. Yang, P. R. Montague, D. M. Eagleman, Vividness of mental imagery: Individual variability can be measured objectively. Vision Res. 47, 474–478 (2007).

A. M. Dale, Optimal experimental design for event-related fMRI. Hum. Brain Mapp. 8, 109–114 (1999).

W. A. de Heer, A. G. Huth, T. L. Griffiths, J. L. Gallant, F. E. Theunissen, The hierarchical cortical organization of human speech processing. J. Neurosci. 37, 6539–6557 (2017).

B. Deen, W. A. Freiwald, Parallel systems for social and spatial reasoning within the cortical apex. BioRxiv 2021–09 (2021).

N. Dijkstra, Uncovering the role of the early visual cortex in visual mental imagery. Vision 8, 29 (2024).

L. M. DiNicola, R. M. Braga, R. L. Buckner, Parallel distributed networks dissociate episodic and social functions within the individual. J. Neurophysiol. 123, 1144–1179 (2020).

L. M. DiNicola, O. I. Ariyo, R. L. Buckner, Functional specialization of parallel distributed networks revealed by analysis of trial-to-trial variation in processing demands. J. Neurophysiol. 129, 17–40 (2023).

G. Doucet, M. Naveau, L. Petit, L. Zago, F. Crivello, G. Jobard, N. Delcroix, E. Mellet, N. Tzourio-Mazoyer, B. Mazoyer, M. Joliot, Patterns of hemodynamic low-frequency oscillations in the brain are modulated by the nature of free thought during rest. Neuroimage 59, 3194–3200 (2012).

J. Du, L. M. DiNicola, P. A. Angeli, N. Saadon-Grosman, W. Sun, S. Kaiser, J. Ladopoulou, A. Xue, B. T. T. Yeo, M. Eldaief, R. L. Buckner, Organization of the human cerebral cortex estimated within individuals: networks, global topography, and function. J. Neurophysiol. 131, 1014–1082 (2024).

D. Edmonds, J. J. Salvo, N. Anderson, M. Lakshman, Q. Yang, K. Kay, C. Zelano, R. M. Braga, The human social cognitive network contains multiple regions within the amygdala. Sci. Adv. 10, eadp0453 (2024).

E. Fedorenko, P. J. Hsieh, A. Nieto-Castañón, S. Whitfield-Gabrieli, N. Kanwisher, New method for fMRI investigations of language: defining ROIs functionally in individual subjects. J. Neurophysiol. 104, 1177–1194 (2010).

E. Fedorenko, A. A. Ivanova, T. I. Regev, The language network as a natural kind within the broader landscape of the human brain. Nat. Rev. Neuro. 25, 1–24 (2024).

D. J. Felleman, D. C. Van Essen, Distributed hierarchical processing in the primate cerebral cortex. Cereb. Cortex 1, 1–47 (1991).

C. Fernyhough, A. M. Borghi, Inner speech as language process and cognitive tool. Trends Cogn. Sci. 27, 1180–1193 (2023).

B. Fischl, M. I. Sereno, A. M. Dale, Cortical surface-based analysis: II: inflation, flattening, and a surface-based coordinate system. Neuroimage 9, 195–207 (1999).

G. Ganis, W. L. Thompson, S. M. Kosslyn, Brain areas underlying visual mental imagery and visual perception: an fMRI study. Cogn. Brain Res. 20, 226–241 (2004).

A. W. Gilmore, A. Quach, S. E. Kalinowski, S. J. Gotts, D. L. Schacter, A. Martin, Dynamic content reactivation supports naturalistic autobiographical recall in humans. J. Neurosci. 41, 153–166 (2021).

M. F. Glasser, T. S. Coalson, E. C. Robinson, C. D. Hacker, J. Harwell, E. Yacoub, K. Ugurbil, J. Andersson, C. F. Beckmann, M. Jenkinson, S. M. Smith, D. C. Van Essen, A multi-modal parcellation of human cortex. Nature 536, 171–178 (2016).

P. S. Goldman-Rakic, Topography of cognition: parallel distributed networks in primate association cortex. Ann. Rev. Neurosci. 11, 137–156 (1988).

E. M. Gordon, T. O. Laumann, A. W. Gilmore, D. J. Newbold, D. J. Greene, J. J. Berg, M. Ortega, C. Hoyt-Drazen, C. Gratton, H. Sun, J. M. Hampton, R. S. Coalson, A. L. Nguyen, K. B. McDermott, J. S. Shimony, A. Z. Snyder, B. L. Schlaggar, S. E. Petersen, S. M. Nelson, N. U. F. Dosenbach, Precision functional mapping of individual human brains. Neuron 95, 791–807 (2017).

C. D. Hacker, T. O. Laumann, N. P. Szrama, A. Baldassarre, A. Z. Snyder, E. C. Leuthardt, M. Corbetta, Resting state network estimation in individual subjects. NeuroImage 82, 616–633 (2013).

D. J. Hagler Jr, S. Hatton, M. D. Cornejo, C. Makowski, D. A. Fair, A. S. Dick, M. T. Sutherland, B. J. Casey, D. M. Barch, M. P. Harms, R. Watts, J. M. Bjork, H. P. Garavan, L. Hilmer, C. J. Pung, C. S. Sicat, J. Kuperman, H. Bartsch, F. Xue, M. M. Heitzeg, A. R. Laird, T. T. Trinh, R. Gonzalez, S. F. Tapert, M. C. Riedel, L. M. Squeglia, L. W. Hyde, M. D. Rosenberg, E. A. Earl, K. D. Howlett, F. C. Baker, M. Soules, J. Diaz, O. Ruiz de Leon, W. K. Thompson, M. C. Neale, M. Herting, E. R. Sowell, R. P. Alvarez, S. W. Hawes, M. Sanchez, J. Bodurka, F. J. Breslin, A. S. Morris, M. P. Paulus, W. K. Simmons, J. R. Polimeni, A. van der Kouwe, A. S. Nencka, K. M. Gray, C. Pierpaoli, J. A. Matochik, A. Noronha, W. M. Aklin, K. Conway, M. Glantz, E. Hoffman, R. Little, M. Lopez, V. Pariyadath, S. R. B. Weiss, D. L. WolN-Hughes, R. DelCarmen-Wiggins, S. W. F. Ewing, O. Miranda-Dominguez, B. J. Nagel, A. J. Perrone, D. T. Sturgeon, A. Goldstone, A. Pfefferbaum, K. M. Pohl, D. Prouty, K. Uban, S. Y. Bookheimer, M. Dapretto, A. Galvan, K. Bagot, J. Giedd, M. A. Infante, J. Jacobus, K. Patrick, P. D. Shilling, R. Desikan, Y. Li, L. Sugrue, M. T. Banich, N. Friedman, J. K. Hewitt, C. Hopfer, J. Sakai, J. Tanabe, L. B. Cottler, S. J. Nixon, L. Chang, C. Cloak, T. Ernst, G. Reeves, D. N. Kennedy, S. Heeringa, S. Peltier, J. Schulenberg, C. Sripada, R. A. Zucker, W. G. Iacono, M. Luciana, F. J. Calabro, D. B. Clark, D. A. Lewis, B. Luna, C. Schirda, T. Brima, J. J. Foxe, E. G. Freedman, D. W. Mruzek, M. J. Mason, R. Huber, E. McGlade, A. Prescot, P. F. Renshaw, D. A. Yurgelun-Todd, N. A. Allgaier, J. A. Dumas, M. Ivanova, A. Potter, P. Florsheim, C. Larson, K. Lisdahl, M. E. Charness, B. Fuemmeler, J. M. Hettema, H. H. Maes, J. Steinberg, A. P. Anokhin, P. Glaser, A. C. Heath, P. A. Madden, A. B. Sommers, R. T. Constable, S. J. Grant, G. J. Dowling, S. A. Brown, T. L. Jernigan, A. M. Dale, Image processing and analysis methods for the Adolescent Brain Cognitive Development Study. Neuroimage 202, 116091 (2019).

A. R. Halpern, R. J. Zatorre, When that tune runs through your head: a PET investigation of auditory imagery for familiar melodies. Cereb. Cortex 9, 697–704 (1999).

P. A. Harris, R. Taylor, R. Thielke, J. Payne, N. Gonzalez, J. G. Conde, Research electronic data capture (REDCap)—a metadata-driven methodology and workflow process for providing translational research informatics support. J. Biomed. Inform. 42, 377–381 (2009).

P. A. Harris, R. Taylor, B. L. Minor, V. Elliott, M. Fernandez, L. O’Neal, L. McLeod, G. Delacqua, F. Delacqua, J. Kirby, S. N. Duda, REDCap Consortium, The REDCap consortium: building an international community of software platform partners. J. Biomed. Inform. 95, 103208 (2019).

D. Hassabis, R. N. Spreng, A. A. Rusu, C. A. Robbins, R. A. Mar, D. L. Schacter, Imagine all the people: how the brain creates and uses personality models to predict behavior. Cereb. Cortex 24, 1979–1987 (2014).

S. C. Herholz, A. R. Halpern, R. J. Zatorre, Neuronal correlates of perception, imagery, and memory for familiar tunes. J. Cogn. Neurosci. 24, 1382–1397 (2012).

S. Heunis, M. Breeuwer, C. Caballero-Gaudes, L. Hellrung, W. Huijbers, J. F. Jansen, R. Lamerichs, S. Zinger, A. P. Aldenkamp, The effects of multi-echo fMRI combination and rapid T2*-mapping on offline and real-time BOLD sensitivity. Neuroimage 238, 118244 (2021).

G. Hickok, D. Poeppel, The cortical organization of speech processing. Nat. Rev. Neurosci. 8, 393–402 (2007).

R. T. Hurlburt, B. Alderson-Day, S. Kühn, C. Fernyhough, Exploring the ecological validity of thinking on demand: Neural correlates of elicited vs. spontaneously occurring inner speech. PLoS One 11, e0147932 (2016).

T. Iwata, T. Yanagisawa, Y. Ikegaya, J. Smallwood, R. Fukuma, S. Oshino, N. Tani, H. M. Khoo, H. Kishima, Hippocampal sharp-wave ripples correlate with periods of naturally occurring self-generated thoughts in humans. Nat. Commun. 15, 4078 (2024).

R. JackendoN, How language helps us think. Pragmatics Cogn. 4, 1–34 (1996).

R. JackendoN, What is the human language faculty? Two views. Language 586–624 (2011).

M. Jenkinson, P. Bannister, M. Brady, S. Smith, Improved optimization for the robust and accurate linear registration and motion correction of brain images. Neuroimage 17, 825–841 (2002).

N. Kanwisher, Functional specificity in the human brain: a window into the functional architecture of the mind. Proc. Natl. Acad. Sci. U.S.A. 107, 11163–11170 (2010).

B. Kleber, N. Birbaumer, R. Veit, T. Trevorrow, M. Lotze, Overt and imagined singing of an Italian aria. Neuroimage 36, 889-900 (2007).

R. Kong, J. Li, C. Orban, M. R. Sabuncu, H. Liu, A. Schaefer, N. Sun, X. Zuo, A. J. Jones, S. B. EickhoN, B. T. T. Yeo, Spatial topography of individual-specific cortical networks predicts human cognition, personality, and emotion. Cereb. Cortex 29, 2533–2551 (2019).

T. Konkle, A. Oliva, Canonical visual size for real-world objects. J. Exp. Psychol. Hum. Percept. Perform. 37, 23 (2011).

D. Konu, B. Mckeown, A. Turnbull, N. S. P. Ho, T. Karapanagiotidis, T. Vanderwal, C. McCall, S. P. Tipper, E. Jefferies, J. Smallwood, Exploring patterns of ongoing thought under naturalistic and conventional task-based conditions. Conscious. Cogn. 93, 103139 (2021).

D. Konu, A. Turnbull, T. Karapanagiotidis, H. T. Wang, L. R. Brown, E. Jefferies, J. Smallwood, A role for the ventromedial prefrontal cortex in self-generated episodic social cognition. Neuroimage 218, 116977 (2020).

S. M. Kosslyn, W. L. Thompson, When is early visual cortex activated during visual mental imagery? Psychol. Bull. 129, 723 (2003).

S. M. Kosslyn, W. L. Thompson, N. M. Alpert, Neural systems shared by visual imagery and visual perception: A positron emission tomography study. Neuroimage 6, 320–334 (1997).

S. M. Kosslyn, W. L. Thompson, G. Ganis. The case for mental imagery. (Oxford University Press, Oxford, UK, 2006).

D. J. Kraemer, C. N. Macrae, A. E. Green, W. M. Kelley, Sound of silence activates auditory cortex. Nature 434, 158 (2005).

L. Krubitzer, The magnificent compromise: cortical field evolution in mammals. Neuron 56, 201–208 (2007).

Y. H. Kwon, J. J. Salvo, N. L. Anderson, D. Edmonds, A. M. Holubecki, M. Lakshman, K. Yoo, B. T. T. Yeo, K. Kay, C. Gratton, R. M. Braga, Situating the salience and parietal memory networks in the context of multiple parallel distributed networks using precision functional mapping. Cell Rep. 44, 1 (2025).

S. Lambert, E. Sampaio, C. Scheiber, Y. Mauss, Neural substrates of animal mental imagery: calcarine sulcus and dorsal pathway involvement—an fMRI study. Brain Res. 924, 176–183 (2002).

T. O. Laumann, E. M. Gordon, B. Adeyemo, A. Z. Snyder, S. J. Joo, M. Y. Chen, A. W. Gilmore, K. B. McDermott, S. M. Nelson, N. U. F. Dosenbach, B. L. Schlaggar, J. A. Mumford, R. A. Poldrack, S. E. Petersen, Functional system and areal organization of a highly sampled individual human brain. Neuron 87, 657–670 (2015).

M. H. Lee, C. D. Hacker, A. Z. Snyder, M. Corbetta, D. Zhang, E. C. Leuthardt, J. S. Shimony, Clustering of resting state networks. PLOS One 7, e40370 (2012).

R. L. Lewis, S. Vasishth, J. A. Van Dyke, Computational principles of working memory in sentence comprehension. Trends Cogn. Sci. 10, 447–454 (2006).

L. Lu, M. Han, G. Zou, L. Zheng, J. H. Gao, Common and distinct neural representations of imagined and perceived speech. Cereb. Cortex 33, 6486–6493 (2023).

C. J. Lynch, I. Elbau, C. Liston, Improving precision functional mapping routines with multi-echo fMRI. Curr. Opin. Behav. Sci. 40, 113–119 (2021).

D. S. Margulies, S. S. Ghosh, A. Goulas, M. Falkiewicz, J. M. Huntenburg, G. Langs, G. Bezgin, S. B. EickhoN, F. X. Castellanos, M. Petrides, E. Jefferies, J. Smallwood, Situating the default-mode network along a principal gradient of macroscale cortical organization. Proc. Natl. Acad. Sci. U.S.A. 113, 12574–12579 (2016).

D. F. Marks, Visual imagery differences in the recall of pictures. Br. J. Psychol. 64, 17–24 (1973).

J. C. Mazziotta, A. W. Toga, A. Evans, P. Fox, J. Lancaster, A probabilistic atlas of the human brain: theory and rationale for its development. Neuroimage 2, 89–101 (1995).

P. K. McGuire, D. A. Silbersweig, R. M. Murray, A. S. David, R. S. J. Frackowiak, C. D. Frith, Functional anatomy of inner speech and auditory verbal imagery. Psychol. Med. 26, 29–38 (1996).

B. Mckeown, W. H. Strawson, M. Zhang, A. Turnbull, D. Konu, T. Karapanagiotidis, H. T. Wang, R. Leech, T. Xu, S. Hardikar, B. Bernhardt, D. Margulies, E. Jefferies, J. Wammes, J. Smallwood, Experience sampling reveals the role that covert goal states play in task-relevant behavior. Sci. Rep. 13, 21710 (2023).

V. Menon, 20 years of the default mode network: A review and synthesis. Neuron 111, 2469–2487 (2023).

M. M. Mesulam, Large-scale neurocognitive networks and distributed processing for attention, language, and memory. Ann. Neurol. 28, 597–613 (1990).

M. M. Mesulam, From sensation to cognition. Brain 121, 1013–1052 (1998).

B. Mulholland, L. Chitiz, R. Wallace, B. Mckeown, M. Milham, A. Klein, R. Leech, E. Jefferies, G. Poerio, J. Wammes, J. Stewart, S. Hardikar, J. Smallwood, Patterns of Ongoing Thought in the Real World and Their Links to Mental Health and Well-Being. biorxiv 2024–07 (2024).

B. Mulholland, I. Goodall-Halliwell, R. Wallace, L. Chitiz, B. Mckeown, A. Rastan, G. Poerio, R. Leech, A. Turnbull, A. Klein, M. Milham, J. Wammes, E. Jefferies, J. Smallwood, Patterns of ongoing thought in the real world. Conscious. Cogn. 114, 103530 (2023).

T. Naselaris, C. A. Olman, D. E. Stansbury, K. Ugurbil, J. L. Gallant, A voxel-wise encoding model for early visual areas decodes mental images of remembered scenes. Neuroimage 105, 215–228 (2015).

K. M. O’Craven, N. Kanwisher, Mental imagery of faces and places activates corresponding stimulus-specific brain regions. J. Cogn. Neurosci. 12, 1013–1023 (2000).

M. Papoutsi, J. A. de Zwart, J. M. Jansma, M. J. Pickering, J. A. Bednar, B. Horwitz, From phonemes to articulatory codes: an fMRI study of the role of Broca’s area in speech production. Cereb. Cortex 19, 2156–2165 (2009).

E. Paulesu, C. D. Frith, R. S. Frackowiak, The neural correlates of the verbal component of working memory. Nature 362, 342–345 (1993).

M. Peer, R. Salomon, I. Goldberg, O. Blanke, S. Arzy, Brain system for mental orientation in space, time, and person. Proc. Natl. Acad. Sci. U.S.A. 112, 11072–11077 (2015).

M. Perrone-Bertolotti, J. Kujala, J. R. Vidal, C. M. Hamame, T. Ossandon, O. Bertrand, L. Minotti, P. Kahane, K. Jerbi, J. P. Lachaux, How silent is silent reading? Intracerebral evidence for top-down activation of temporal voice areas during reading. J. Neurosci. 32, 17554–17562 (2012).

B. A. Poser, M. J. Versluis, J. M. Hoogduin, D. G. Norris, BOLD contrast sensitivity enhancement and artifact reduction with multiecho EPI: parallel-acquired inhomogeneity-desensitized fMRI. Magn. Reson. Med. 55, 1227–1235 (2006).

J. Pratts, G. Pobric, B. Yao, Bridging phenomenology and neural mechanisms of inner speech: ALE meta-analysis on egocentricity and spontaneity in a dual-mechanistic framework. Neuroimage 282, 120399 (2023).

R Core Team, R: A language and environment for statistical computing. R Found. Stat. Comput. Vienna, Austria (2021).

M. E. Raichle, A. M. MacLeod, A. Z. Snyder, W. J. Powers, D. A. Gusnard, G. L. Shulman, A default mode of brain function. Proc. Natl. Acad. Sci. U.S.A. 98, 676–682 (2001).

K. Rastle, J. Harrington, M. Coltheart, 358,534 nonwords: The ARC nonword database. Q. J. Exp. Psychol. A 55, 1339–1362 (2002).

J. J. Salvo, N. L. Anderson, R. M. Braga, Intrinsic functional connectivity delineates transmodal language functions. bioRxiv 2024–12 (2024).

R. Saxe, Uniquely human social cognition. Curr. Opin. Neurobiol. 16, 235–239 (2006).

Z. M. Saygin, D. E. Osher, E. S. Norton, D. A. Youssoufian, S. D. Beach, J. Feather, N. Gaab, J. D. E. Gabrieli, N. Kanwisher, Connectivity precedes function in the development of the visual word form area. Nat. Neurosci. 19, 1250–1255 (2016).

D. L. Schacter, D. R. Addis, On the nature of medial temporal lobe contributions to the constructive simulation of future events. Philos. Trans. R. Soc. B 364, 1245–1253 (2009).

D. L. Schacter, D. R. Addis, R. L. Buckner, Remembering the past to imagine the future: The prospective brain. Nat. Rev. Neurosci. 8, 657–661 (2007).

T. L. Scott, J. Gallée, E. Fedorenko, A new fun and robust version of an fMRI localizer for the frontotemporal language system. Cogn. Neurosci. 8, 167–176 (2017).

S. S. Shergill, E. T. Bullmore, M. J. Brammer, S. C. R. Williams, R. M. Murray, P. K. McGuire, A functional study of auditory verbal imagery. Psychol. Med. 31, 241–253 (2001).

S. S. Shergill, M. J. Brammer, R. Fukuda, E. Bullmore, E. Amaro Jr, R. M. Murray, P. K. McGuire, Modulation of activity in temporal cortex during generation of inner speech. Hum. Brain Mapp. 16, 219–227 (2002).

G. L. Shulman, M. Corbetta, R. L. Buckner, J. A. Fiez, F. M. Miezin, M. E. Raichle, S. E. Petersen, Common blood flow changes across visual tasks: I. Increases in subcortical structures and cerebellum but not in nonvisual cortex. J. Cogn. Neurosci. 9, 624–647 (1997).

E. H. Silson, A. Steel, A. Kidder, A. W. Gilmore, C. I. Baker, Distinct subdivisions of human medial parietal cortex are recruited differentially for memory recall of places and people. bioRxiv 554915 (2019a).

E. H. Silson, A. W. Gilmore, S. E. Kalinowski, A. Steel, A. Kidder, A. Martin, C. I. Baker, A posterior-anterior distinction between scene perception and scene construction in human medial parietal cortex. J. Neurosci. 39, 705–717 (2019b).

S. D. Slotnick, W. L. Thompson, S. M. Kosslyn, Visual mental imagery induces retinotopically organized activation of early visual areas. Cereb. Cortex 15, 1570–1583 (2005).

J. Smallwood, L. Nind, R. C. O’Connor, When is your head at? An exploration of the factors associated with the temporal focus of the wandering mind. Conscious. Cogn. 18, 118–125 (2009).

J. Smallwood, T. Karapanagiotidis, F. Ruby, B. Medea, I. De Caso, M. Konishi, H. T. Wang, G. Hallam, D. S. Margulies, E. Jefferies, Representing representation: Integration between the temporal lobe and the posterior cingulate influences the content and form of spontaneous thought. PLoS One 11, e0152272 (2016).

J. Smallwood, A. Turnbull, H. T. Wang, N. S. Ho, G. L. Poerio, T. Karapanagiotidis, D. Konu, B. Mckeown, M. Zhang, C. Murphy, D. Vatansever, D. Bzdok, M. Konishi, R. Leech, P. Seli, J. W. Schooler, B. Bernhardt, D. S. Margulies, E. Jefferies, The neural correlates of ongoing conscious thought. iScience 24, 3 (2021).

R. N. Spreng, J. R. Andrews-Hanna, The default network and social cognition. Brain Mapping 3, 165–169 (2015).

A. Spagna, D. Hajhajate, J. Liu, P. Bartolomeo, Visual mental imagery engages the left fusiform gyrus, but not the early visual cortex: A meta-analysis of neuroimaging evidence. Neurosci. Biobehav. Rev. 122, 201–217 (2021).

R. N. Spreng, C. L. Grady, Patterns of brain activity supporting autobiographical memory, prospection, and theory of mind, and their relationship to the default mode network. J. Cogn. Neurosci. 22, 1112–1123 (2010).

R. N. Spreng, W. D. Stevens, J. P. Chamberlain, A. W. Gilmore, D. L. Schacter, Default network activity, coupled with the frontoparietal control network, supports goal-directed cognition. NeuroImage 53, 303–317 (2010).

A. Steel, M. M. Billings, E. H. Silson, C. E. Robertson, A network linking scene perception and spatial memory systems in posterior cerebral cortex. Nat. Commun. 12, 2632 (2021).

A. Steel, B. D. Garcia, K. Goyal, A. Mynick, C. E. Robertson, Scene perception and visuospatial memory converge at the anterior edge of visually responsive cortex. J. Neurosci. 43, 5723–5737 (2023).

A. Steel, B. D. Garcia, D. Prasad, C. E. Robertson, Topography of scene memory and perception activity in posterior cortex–a publicly available resource. bioRxiv 2025–01 (2025).

M. Stephane, M. Dzemidzic, G. Yoon, Keeping the inner voice inside the head, a pilot fMRI study. Brain Behav. 11, e02042 (2021).

A. A. Sulfaro, A. K. Robinson, T. A. Carlson, Properties of imagined experience across visual, auditory, and other sensory modalities. Conscious. Cogn. 117, 103598 (2024).

X. Tian, J. M. Zarate, D. Poeppel, Mental imagery of speech implicates two mechanisms of perceptual reactivation. Cortex 77, 1–12 (2016).

M. D. Tisdall, A. T. Hess, M. Reuter, E. M. Meintjes, B. Fischl, A. J. van der Kouwe, Volumetric navigators for prospective motion correction and selective reacquisition in neuroanatomical MRI. Magn. Reson. Med. 68, 389–399 (2012).

A. Turnbull, H. T. Wang, C. Murphy, N. S. P. Ho, X. Wang, M. Sormaz, T. Karapanagiotidis, R. M. Leech, B. Bernhardt, D. S. Margulies, D. Vatansever, E. Jefferies, J. Smallwood, Left dorsolateral prefrontal cortex supports context-dependent prioritisation of oN-task thought. Nat. Commun. 10, 3816 (2019).

R. S. Varrier, E. S. Finn, Seeing social: A neural signature for conscious perception of social interactions. J. Neurosci. 42, 9211–9226 (2022).

D. Vatansever, D. Bzdok, H. T. Wang, G. Mollo, M. Sormaz, C. Murphy, T. Karapanagiotidis, J. Smallwood, E. Jefferies, Varieties of semantic cognition revelated through simultaneous decomposition of intrinsic brain connectivity and behavior. Neuroimage 158, 1–11 (2017).

M. Villena-Gonzalez, H. T. Wang, M. Sormaz, G. Mollo, D. S. Margulies, E. A. Jefferies, J. Smallwood, Individual variation in the propensity for prospective thought is associated with functional integration between visual and retrosplenial cortex. Cortex 99, 224–234 (2018).

R. S. Wallace, B. Mckeown, I. Goodall-Halliwell, L. Chitiz, P. Forest, T. Karapanagiotidis, B. Mullholand, A. G. Turnbull, T. Vanderwal, S. Hardikar, T. Gonzalez Alam, B. Bernhardt, H. T. Wang, W. Strawson, M. Milham, T. Xu, D. S. Margulies, G. L. Poerio, E. Jefferies, J. I. Skipper, J. D. Wammes, R. Leech, J. Smallwood, Mapping patterns of thought onto brain activity during movie-watching. eLife 13, RP97731 (2025).

H. Wang, N. S. P. Ho, D. Bzdok, B. C. Bernhardt, D. S. Margulies, E. Jefferies, J. Smallwood, Neurocognitive patterns dissociating semantic processing from executive control are linked to more detailed off-task mental time travel. Sci. Rep. 10, 11904 (2020).

C. I. Winlove, F. Milton, J. Ranson, J. Fulford, M. MacKisack, F. Macpherson, A. Zeman, The neural correlates of visual imagery: A co-ordinate-based meta-analysis. Cortex 105, 4–25 (2018).

M. W. Woolrich, B. D. Ripley, M. Brady, S. M. Smith, Temporal autocorrelation in univariate linear modeling of FMRI data. NeuroImage 14, 1370–1386 (2001).

B. Yao, P. Belin, C. Scheepers, Silent reading of direct versus indirect speech activates voice-selective areas in the auditory cortex. J. Cogn. Neurosci. 23, 3146–3152 (2011).

T. Yarkoni, D. M. Barch, J. R. Gray, T. E. Conturo, T. S. Braver, BOLD correlates of trial-by-trial reaction time variability in gray and white matter: A multi-study fMRI analysis. PLoS One 4, e4257 (2009).

B. T. T. Yeo, F. M. Krienen, J. Sepulcre, M. R. Sabuncu, D. Lashkari, M. Hollinshead, J. L. Roffman, J. W. Smoller, L. Zöllei, J. R. Polimeni, B. Fischl, H. Liu, R. L. Buckner, The organization of the human cerebral cortex estimated by intrinsic functional connectivity. J. Neurophysiol. (2011).

S. S. Yoo, C. U. Lee, B. G. Choi, Human brain mapping of auditory imagery: Event-related functional MRI study. NeuroReport 12, 3045–3049 (2001).

A. Zeman, Aphantasia and hyperphantasia: Exploring imagery vividness extremes. Trends Cogn. Sci. (2024).

B. Yuan, H. Xie, Z. Wang, Y. Xu, H. Zhang, J. Liu, L. Chen, C. Li, S. Tan, Z. Lin, X. Hu, T. Gu, J. Lu, D. Liu, J. Wu, The domain-separation language network dynamics in resting state support its flexible functional segregation and integration during language and speech processing. NeuroImage 274, 120132 (2023).

A. Zeman, M. Dewar, S. Della Sala, Lives without imagery–Congenital aphantasia. Cortex 73, 378–380 (2015).

M. Zhang, B. C. Bernhardt, X. Wang, D. Varga, K. Krieger-Redwood, J. Royer, R. Rodríguez-Cruces, R. Vos de Wael, D. S. Margulies, J. Smallwood, E. Jefferies, Perceptual coupling and decoupling of the default mode network during mind-wandering and reading. eLife 11, e74011 (2022).

