## Supplementary Materials for "Distinct distributed brain networks dissociate self-generated mental states"

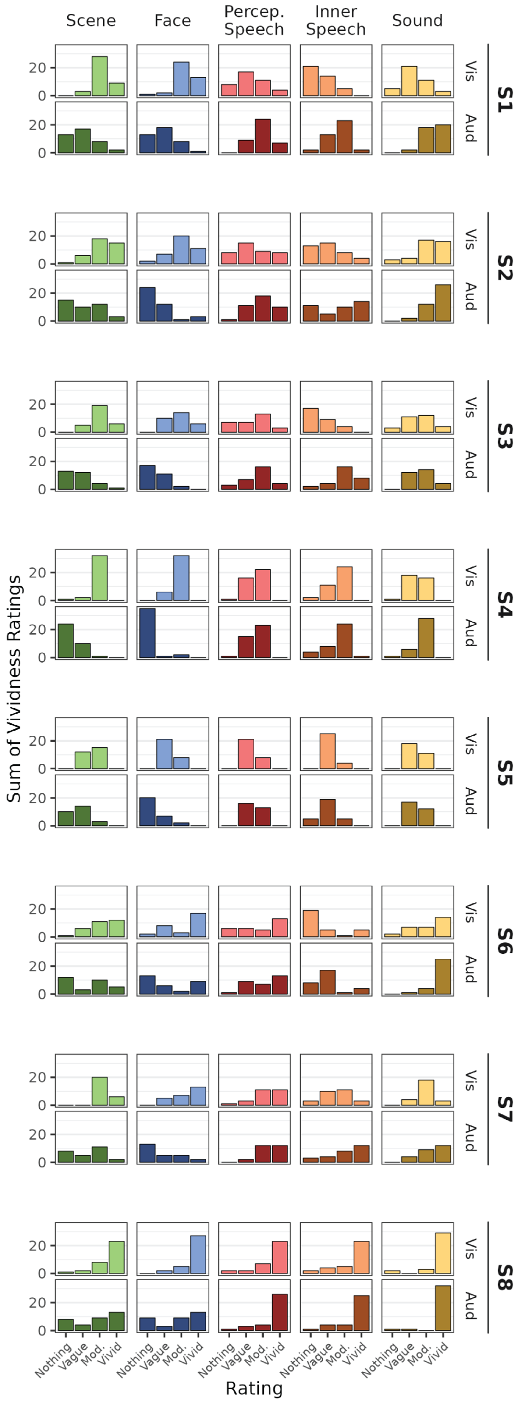


**Supp. Fig. S1: Individual participants' ratings of visual and auditory vividness for each imagination condition.** Breaking down Figure 1C by participant, there is a pattern shared across participants, where scenes and faces tend to be rated higher on visual vividness compared with auditory vividness, while the reverse is true for Perceptual Speech, Inner Speech, and Sound. However, some subjects (e.g. see S8) varied in their response pattern and range of scores utilized.

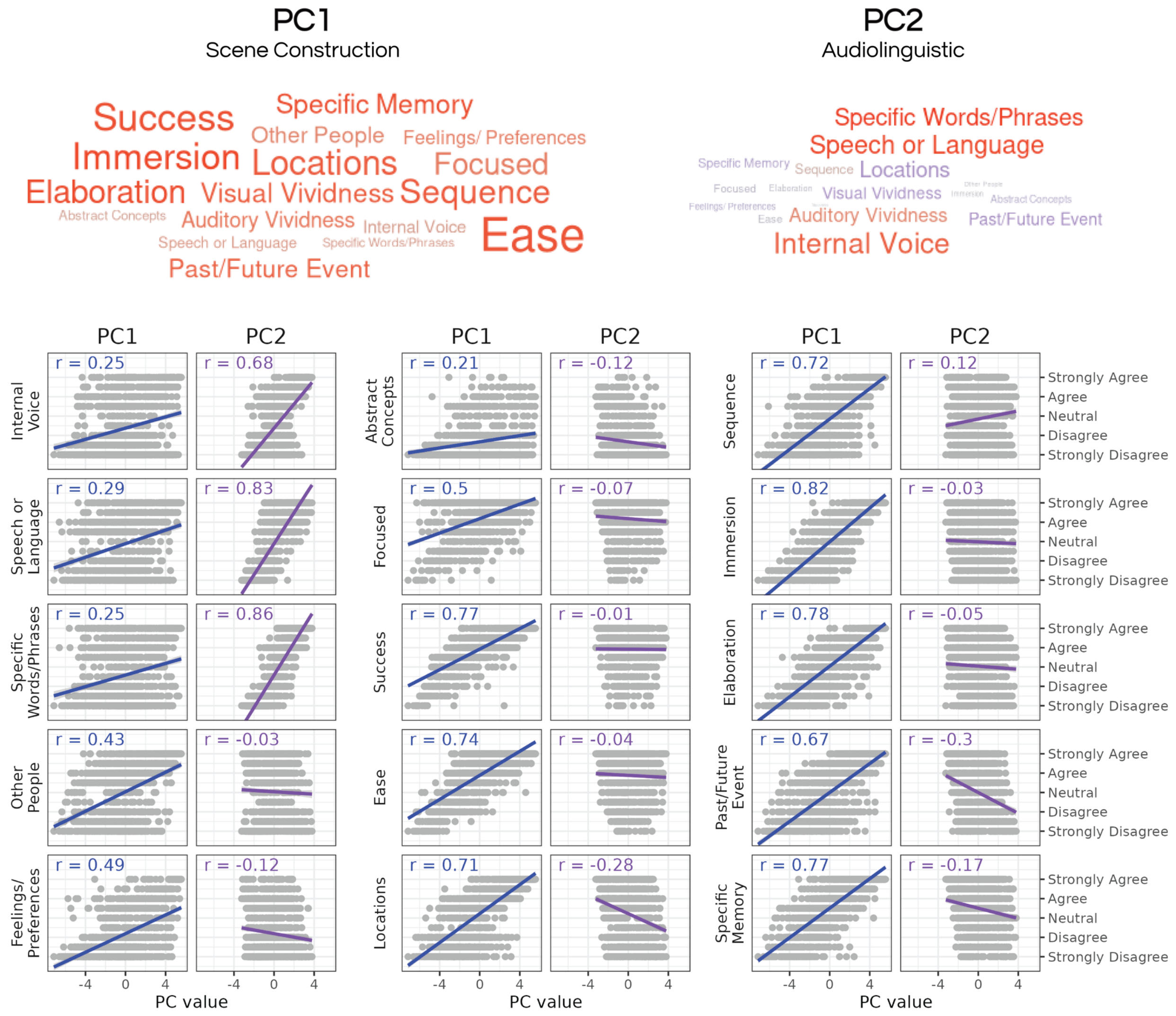


**Supp. Fig. S2: Principal component analysis reveals the interrelationships of the different thought probes.** Top) A word cloud demonstrating the relationship between each probe and the first two principal components, PC1 and PC2. Positive correlations are red, while negative correlations are blue; the size of the text corresponds to the strength of the correlation. Bottom) The correlation is shown between each of the follow-up probes and PC1 (blue) and PC2 (purple). Generally, PC1 shows a positive relationship with all probes, while PC2 shows positive correlations with linguistic probes and negative correlations with probes regarding spatial location and memory.

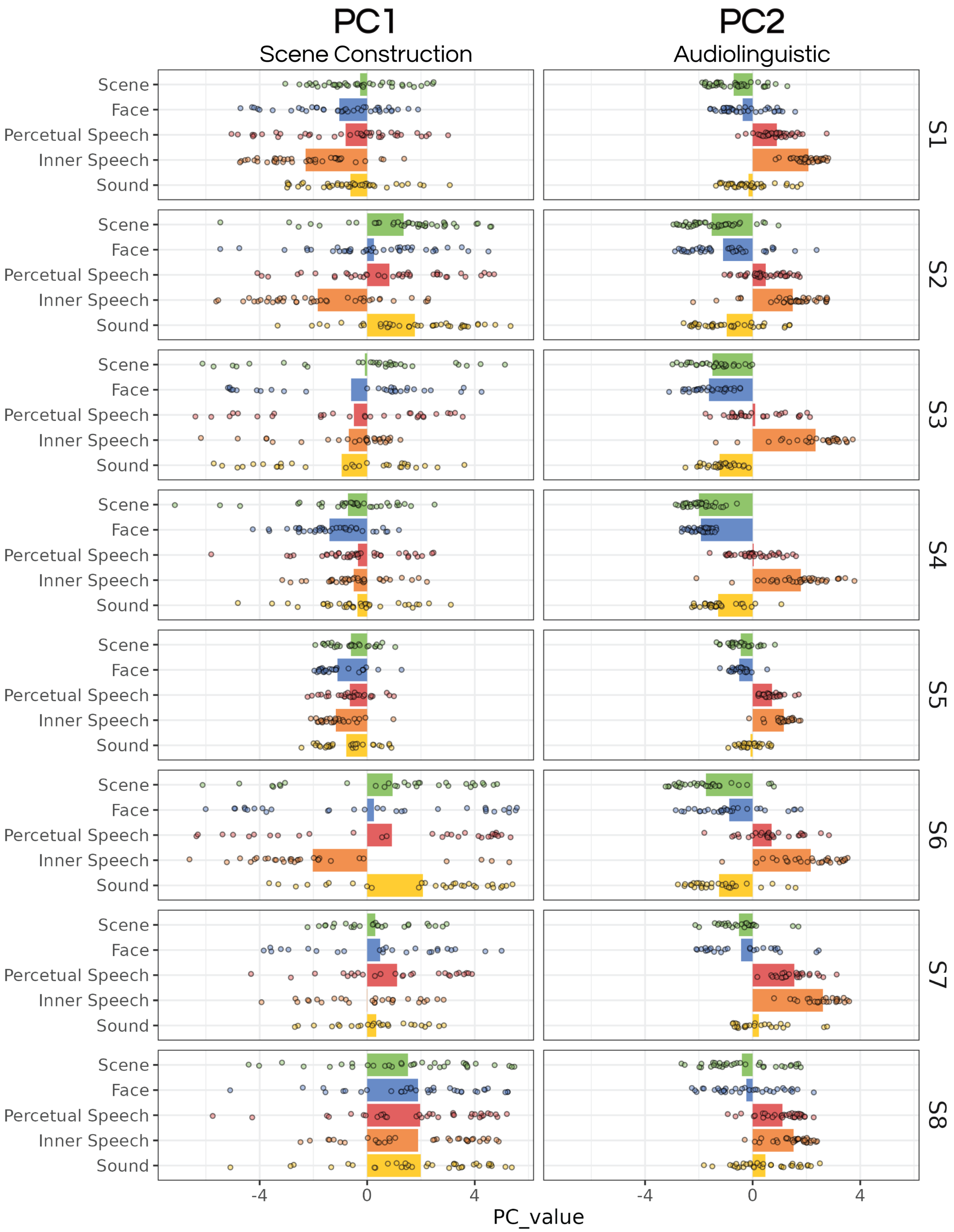


**Supp. Fig. S3: PC1 and PC2 loadings differed across the IMAGINE task conditions.** For the first two principal components from the PCA, the PC value for each trial is plotted along with the averages by stimulus condition (broken down by subject). Subjects show no clear pattern regarding PC1, but generally show more positive PC2 values for the Perceptual Speech and Inner Speech items, and more negative values for Scenes and Faces.

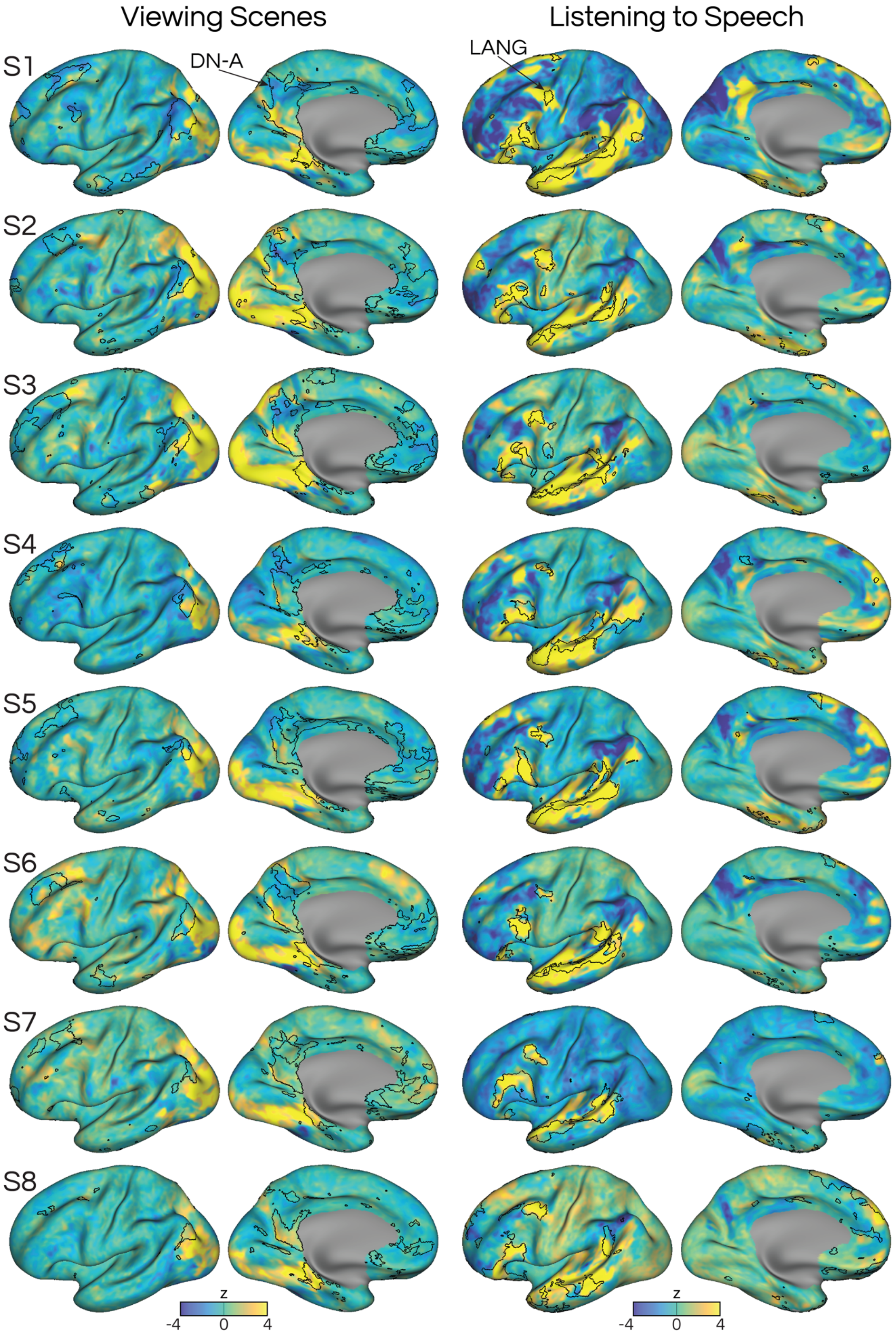
Viewing Scenes
Listening to Speech
S1
DN-A
LANG
S2
S3
S4
S5
S6
S7
S8
z
-4
0
4
z
-4
0
4

**Supp. Fig. 4: Networks DN-A and LANG overlap with activity evoked by viewing scenes and listening to speech.** The left column shows activity when viewing scenes. The right column shows activity when listening to speech, which overlaps largely with the LANG network. The maps in the right and left columns were binarized to create the green regions in Figure 7 and Figure 8, respectively.

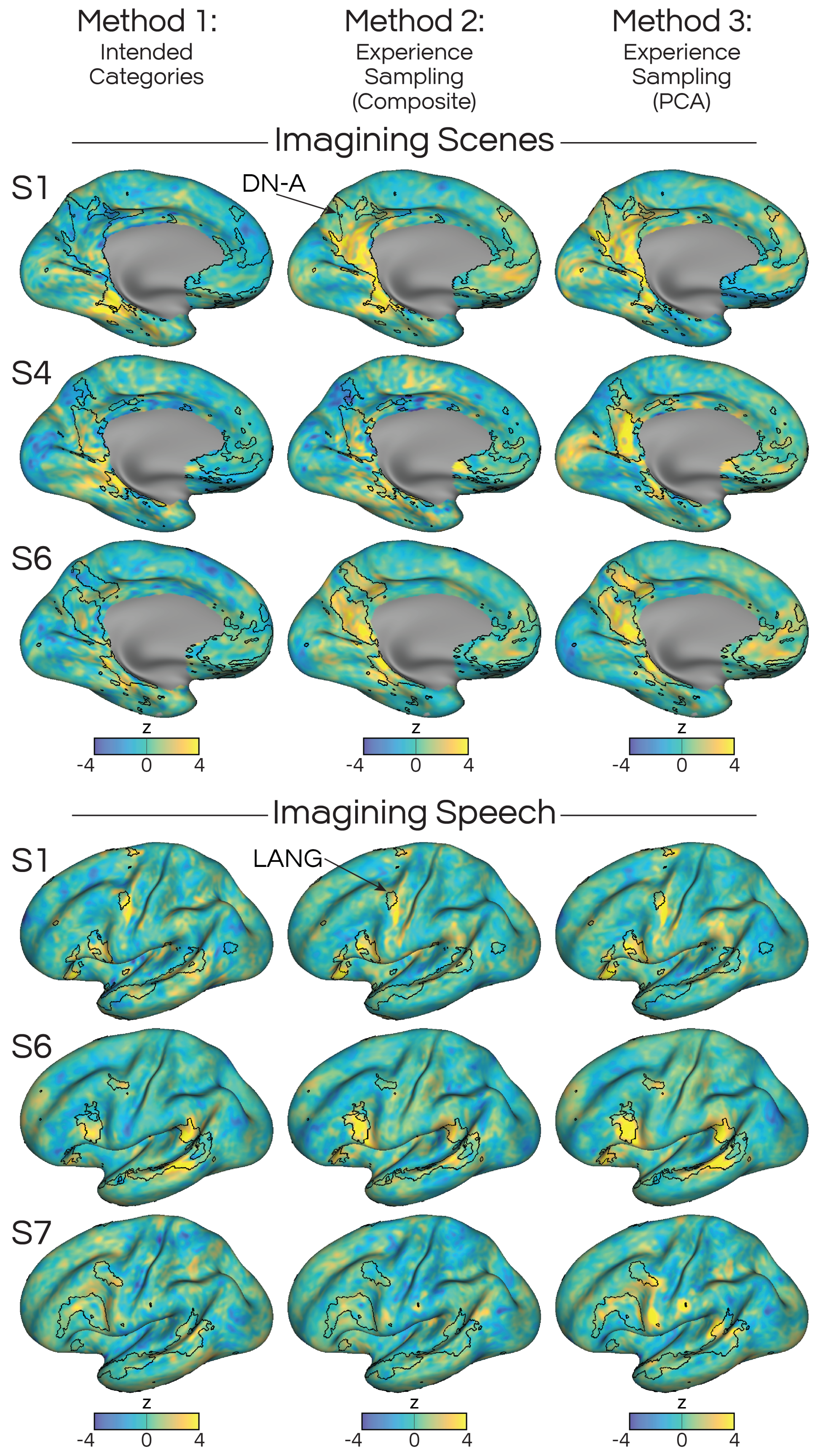
Method 1:
Intended Categories
Method 2:
Experience Sampling (Composite)
Method 3:
Experience Sampling (PCA)
Imagining Scenes
S1
DN-A
S4
S6
z
-4
0
4
Imagining Speech
S1
LANG
S6
S7
z
-4
0
4

**Supp. Fig. S5: Incorporating participants' experience sampling responses improves maps of activity evoked by imagining**. Examples are given for subjects where accounting for participants' trial-wise self-reported ratings about their imagined content improved their task activity maps. In the first column, the *a priori* task conditions were used, and subject ratings were ignored. In the second column, a composite value was created using pre-selected mDES survey ratings, and trials scoring in the top and bottom 20% were contrasted. In the third column, all mDES survey ratings were subjected to a PCA, and the trial-wise loadings for the two principal components, PC1 and PC2, were used as regressors to agglomerate trial-related activity patterns. In these examples, more distributed responses can be seen in the right column, indicating greater sensitivity by accounting for trial-wise variation.

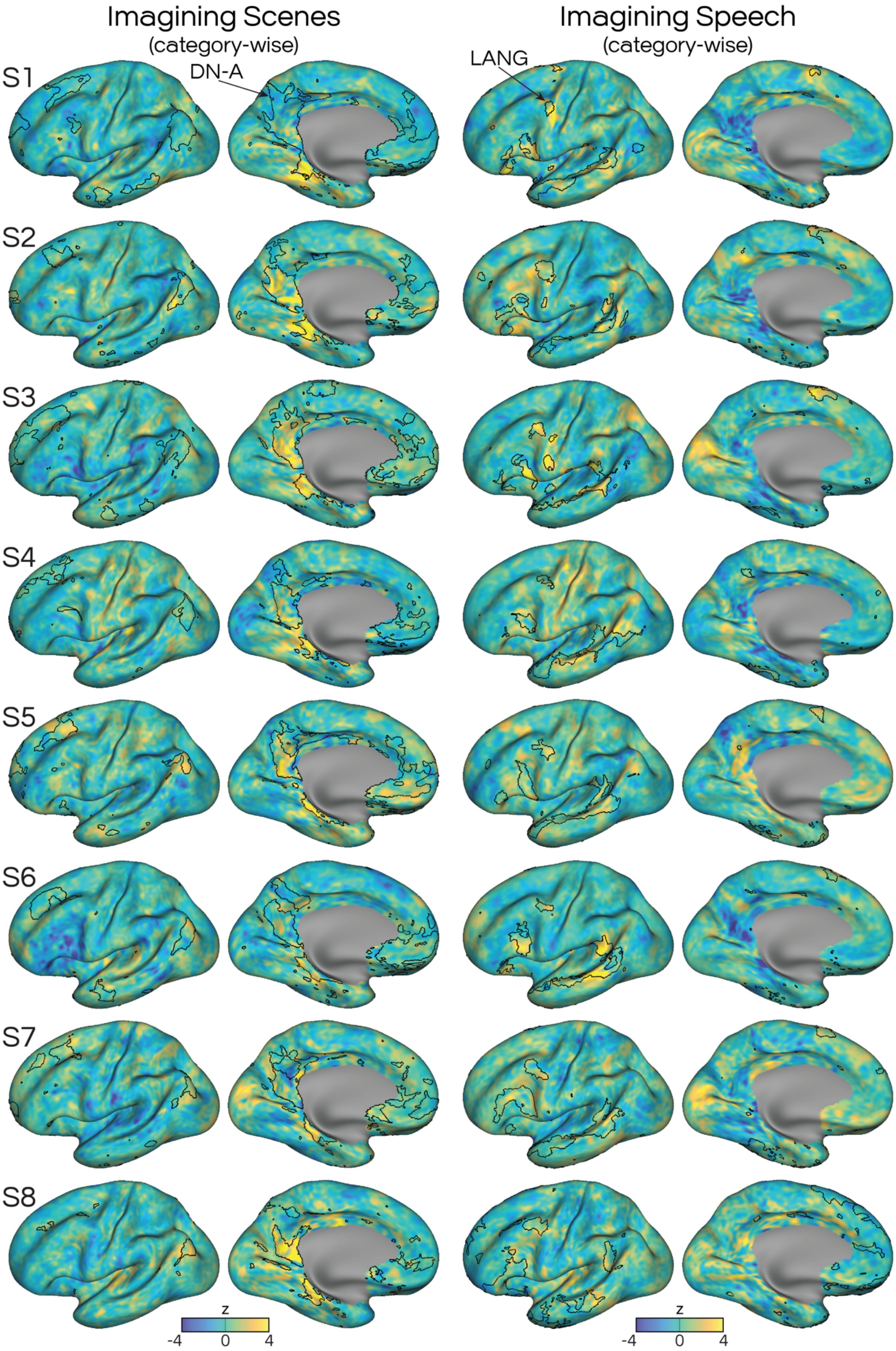
Imagining Scenes
(category-wise)
Imagining Speech
(category-wise)
DN-A
LANG
S1
S2
S3
S4
S5
S6
S7
S8
z
-4
0
4
z
-4
0
4

**Supp. Fig. S6: Condition-wise method for assessing activity evoked during imagining.** The left column shows activity maps for imagining scenes, while the right shows imagining language. These maps were created using condition-wise contrasts (Scenes > Sounds, in the left column, or Perceptual Speech + Inner Speech > Sounds, in the right column). For imagining scenes (left), all participants displayed activity in the parahippocampal region at or near DN-A, but responses elsewhere were inconsistent across participants. For imagining speech (right), two subjects (S3, S6) showed activity overlapping the LANG network, but other results were inconsistent using this condition-wise analysis.

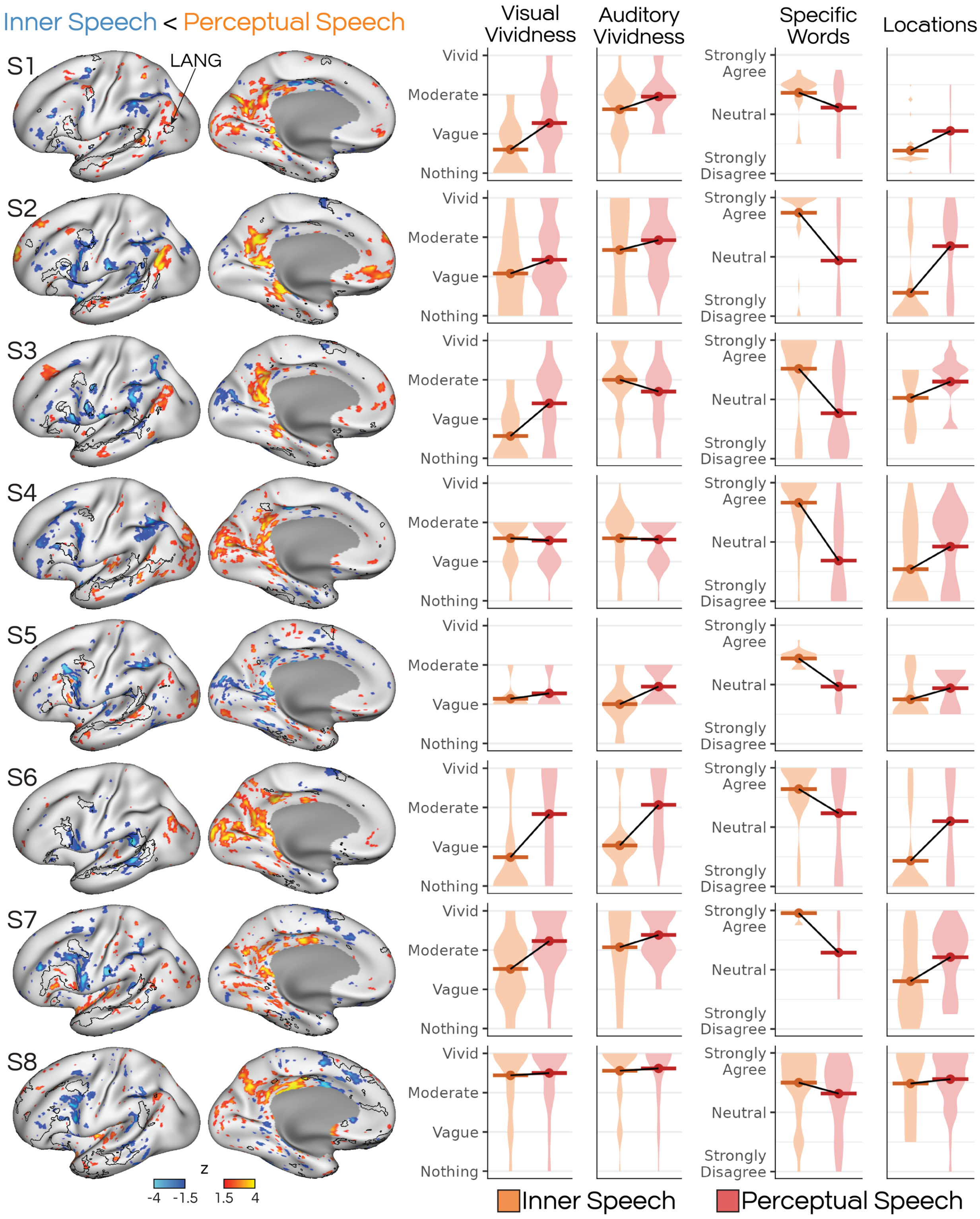

Inner Speech < Perceptual Speech
LANG
S1
S2
S3
S4
S5
S6
S7
S8
z
-4 -1.5 1.5 4
Visual Vividness
Auditory Vividness
Specific Words
Locations
Vivid
Moderate
Vague
Nothing
Strongly Agree
Neutral
Strongly Disagree
Inner Speech
Perceptual Speech

**Supp. Fig. S7: Different forms of imagining speech produce different patterns of brain activity.** Left) Imagined states elicited by the Perceptual Speech condition, which prompted participants to imagine hearing speech, showed greater activity in posteromedial regions (warm colors) than the Inner Speech condition, which prompted participants to imagine specific words. Conversely, imagined states elicited by the Inner Speech condition showed greater activity in frontal and lateral parietal regions (cool colors) that were close to the LANG network and ventral motor strip which controls the orofacial musculature. These results varied across participants. Right) Violin plots show distributions of mDES survey ratings for trials in the Inner Speech and Perceptual Speech conditions. Horizontal bars indicate the means. Mean ratings of visual vividness were generally higher for Perceptual Speech (red) than Inner Speech trials (orange); auditory vividness shows no clear pattern. Ratings for imagining Specific Words were higher for Inner Speech, while ratings for Locations were higher for Perceptual Speech. Thus, imagining different types of speech (inner hearing vs. inner speaking) was related to different mental state properties, and different patterns of brain activity.

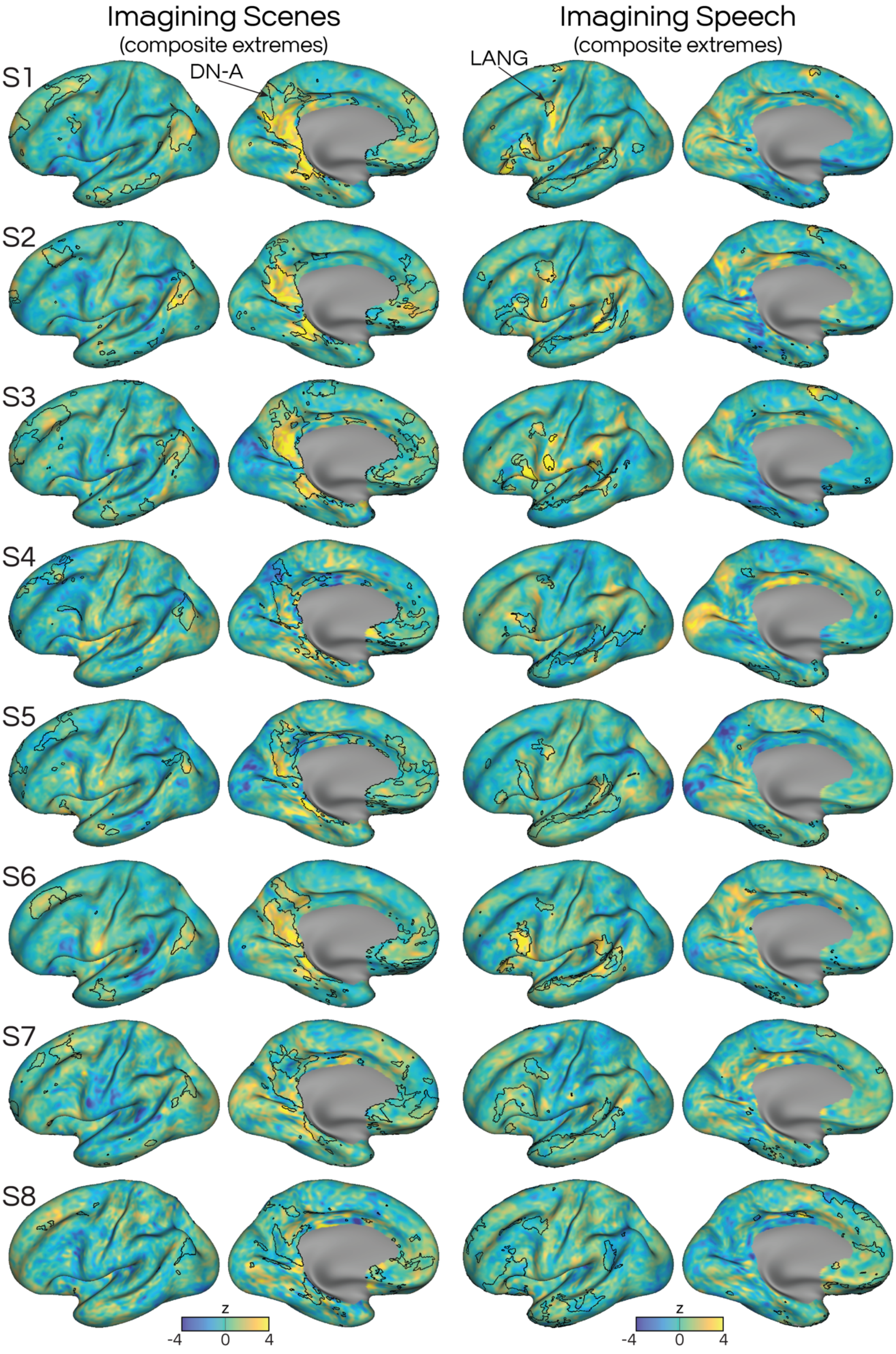

Imagining Scenes
(composite extremes)
Imagining Speech
(composite extremes)
DN-A
LANG
S1
S2
S3
S4
S5
S6
S7
S8
z
-4
0
4
z
-4
0
4

**Supp. Fig. S8: Composite-score method for assessing activity evoked by imagining.** The left column shows activity maps for imagining scenes, while the right shows imagining language. The maps were created using a contrast of the top 20% of items on a composite score, contrasted against the bottom 20% of items on that score. The Imagining Scenes composite included the visual vividness + locations ratings, and the Imagining Speech composite included the auditory vividness + speech/language + specific words/phrases ratings.

**Supplementary Text**

**Instructions given to participants for in-scanner IMAGINE task**

In this task, we're going to ask you to imagine different things while lying still in the scanner. First, **a short sentence will appear on the screen for you to read**, for example: "Imagine a castle on a hill" or "Imagine the sound of a motorcycle engine". Once you have read the sentence, we want you to **follow the instructions**, while remembering to lie very still. You will have seven (7) seconds to read the sentence and think about it. After this, you will be asked to **respond to two questions about what you imagined** (as explained below).

There is no right or wrong way to think about what the sentence instructs; we just want you to think about (imagine) the person, place or thing in whatever way you would normally do that. Similarly, if you are not sure what the instruction is asking you to imagine, that's ok; just try to do the best you can.

**Remember to lie completely still throughout the task** even while you are performing the instructions. For example, if the instruction asks you to "Imagine waving to someone", you don't need to actually wave your arms, just mentally picture yourself doing it. Similarly, if the instructions ask you to "Imagine calling out to a friend", you don't need to actually call out or speak, just think about doing it in your head. **It's especially important not to mouth or move your lips when thinking about speaking**. Instead, just lay completely still and try not to move any muscles.

Sometimes when people perform this kind of imagining, people experience mental sensations, similar to dreaming. For example, if you are asked to "Imagine a red sports car", you might experience a mental visual image of a red sports car, almost as if you were "seeing" it inside your mind. Similarly, if you were asked to "Imagine singing When the Saints Go Marching In", while you do this you might feel as if you are experiencing sounds in your mind, almost as if you were "hearing" the song play internally. These mental sensations are perfectly normal and are the topic of this study.

Sometimes these mental sensations can be really **vivid and clear**, almost like seeing or hearing the item in real life. Sometimes, these mental sensations are **dim or vague**. For example, the mental image may seem dull, or like it is in black and white, or you might not really be able to make out the sounds properly, even though you are still confident that you "saw" or "heard" something. Other times, **you may not experience any mental sensations at all**, but you just 'know' that you are thinking about the item. There is no correct or incorrect way of doing it, and any of the above options may occur depending on what you are thinking about or how you normally think.

Once you have finished the thinking period, we will ask you two questions about whether you experienced these types of mental sensations while following the instructions. Both questions will be presented on the screen, and the question text will show as black when it's time to answer that question.

The first question is **“while imagining this item, how vividly did you experience mental images?”**

For each of the instructions, we want you to **rate the vividness or clarity of your mental sensations on a scale of zero to three**:

- “**Nothing”** means you did not experience any mental images or sounds at all.
- “**Vague**” means you think you did experience mental images or sounds, but they were dim or dull or not very clear.
- “**Moderate**” means you did experience mental images or sounds, and you feel that they were relatively clear or vivid.
- “**Vivid**” means you did experience mental images or sounds, and they were very clear and vivid, almost similar to if you were actually hearing or seeing the item in real life.

The second is **“while imagining this item, how vividly did you experience mental sounds?”**

You will have three (3) seconds to answer each question, but try to answer it as quickly as you can.

It’s important to remember that there is no correct or incorrect answer to any of the questions. We just want you to tell us as honestly as you can whether you experienced these types of mental images or sounds during the thinking period.

After you are done answering the two questions, there will be a pause and then the computer will move on to the next sentence for you to think about. There will be 25 sentences in total.

Do you have any questions?

**Supplementary Table S1: Imagining prompts.** 200 prompts were presented to each participant during fMRI scanning. Prompts were distributed across 5 categories (Scenes, Faces, Perceptual Speech, Inner Speech, and Sounds), with 40 prompts per category.

| Prompt | Category |
|---|---|
| Imagine an art studio | Scene |
| Imagine an old church | Scene |
| Imagine a casino with slot machines | Scene |
| Imagine a children's playground | Scene |
| Imagine a theatrical play on stage | Scene |
| Imagine a movie theater | Scene |
| Imagine an aquarium | Scene |
| Imagine a garden filled with plants | Scene |
| Imagine a back alley with trash bins | Scene |
| Imagine a canyon in a desert | Scene |
| Imagine a nature trail in the summer | Scene |
| Imagine a kite flying high in the sky | Scene |
| Imagine a wall covered with graffiti | Scene |
| Imagine an underpass under a bridge | Scene |
| Imagine the inside of a doctor's office | Scene |
| Imagine a cubicle office with a desk | Scene |
| Imagine a circus performance in a tent | Scene |
| Imagine a thunderstorm over a field | Scene |
| Imagine watching a ballet performance | Scene |
| Imagine a college campus | Scene |
| Imagine a mountain range | Scene |
| Imagine visiting a new museum | Scene |
| Imagine a sunrise over the sea | Scene |
| Imagine an archery range | Scene |
| Imagine walking in a field with tall grass | Scene |
| Imagine a children's birthday party | Scene |
| Imagine a group of ducks swimming | Scene |
| Imagine a public golf course | Scene |
| Imagine walking inside a dark cave | Scene |
| Imagine an ice cream store | Scene |
| Imagine moving through a crowded airport | Scene |
| Imagine a sailboat out on the ocean | Scene |
| Imagine a river running through a forest | Scene |
| Imagine visiting a library filled with books | Scene |
| Imagine a city skyline | Scene |
| Imagine a busy shopping mall | Scene |
| Imagine attending a holiday event | Scene |
| Imagine a swimming pool | Scene |
| Imagine sitting in traffic on the highway | Scene |
| Imagine a high-end jewelry store | Scene |
| Imagine a woman lifting her eyebrow | Face |
| Imagine a clown's face | Face |
| Imagine a person who is afraid | Face |
| Imagine a face wearing makeup | Face |
| Imagine a boy with freckles | Face |
| Imagine the face of a bald man | Face |
| Imagine a woman squinting in the sunlight | Face |
| Imagine a person with a forehead scar | Face |
| Imagine a man with a face tattoo | Face |
| Imagine a boy wearing goggles | Face |
| Imagine the face of a girl with pigtails | Face |
| Imagine someone putting on a face mask | Face |
| Imagine a face with shaving cream | Face |

| Prompt | Category |
|---|---|
| Imagine a teenager's face with acne | Face |
| Imagine a woman with brown eyes | Face |
| Imagine a smiling child with dimples | Face |
| Imagine a girl with braces | Face |
| Imagine a man with an unshaven stubbly chin | Face |
| Imagine a person who is disgusted | Face |
| Imagine the face of an old person | Face |
| Imagine a boy winking | Face |
| Imagine someone with a nose ring | Face |
| Imagine a person who is grimacing in pain | Face |
| Imagine a young girl who is wearing glasses | Face |
| Imagine a person who is confused | Face |
| Imagine a person eating something sour | Face |
| Imagine a criminal's mugshot | Face |
| Imagine a person who is shocked and surprised | Face |
| Imagine a woman laughing | Face |
| Imagine a girl sticking her tongue out | Face |
| Imagine a young man's sunburned face | Face |
| Imagine a woman who is smiling | Face |
| Imagine a man with a mustache | Face |
| Imagine drawing a face with a pencil | Face |
| Imagine a person who is angry | Face |
| Imagine a girl crossing her eyes | Face |
| Imagine a bearded face | Face |
| Imagine a baby's face | Face |
| Imagine a man who is sad | Face |
| Imagine a woman with fake eyelashes | Face |
| Imagine a radio sports announcer | Perceptual Speech |
| Imagine a person rapping | Perceptual Speech |
| Imagine a person with a lisp | Perceptual Speech |
| Imagine a voiceover in a documentary | Perceptual Speech |
| Imagine hearing a crowd at a football game | Perceptual Speech |
| Imagine using a walkie talkie | Perceptual Speech |
| Imagine people talking at a meal | Perceptual Speech |
| Imagine a heavy smoker speaking | Perceptual Speech |
| Imagine someone giving a speech | Perceptual Speech |
| Imagine a person speaking while crying | Perceptual Speech |
| Imagine a person with an accent | Perceptual Speech |
| Imagine a radio interview | Perceptual Speech |
| Imagine a child whining about a toy | Perceptual Speech |
| Imagine a person with a stutter | Perceptual Speech |
| Imagine a radio news story | Perceptual Speech |
| Imagine two people whispering | Perceptual Speech |
| Imagine a person singing loudly | Perceptual Speech |
| Imagine talking on the phone | Perceptual Speech |
| Imagine hearing a couple arguing | Perceptual Speech |
| Imagine a person speaking excitedly | Perceptual Speech |
| Imagine a teacher reading a kid's story | Perceptual Speech |
| Imagine a PA announcement | Perceptual Speech |
| Imagine hearing a teacher talk | Perceptual Speech |
| Imagine someone using baby talk | Perceptual Speech |
| Imagine hearing your voice echoing | Perceptual Speech |
| Imagine a cheer squad chanting | Perceptual Speech |
| Imagine an elevator voice saying the floor | Perceptual Speech |
| Imagine talking to a cashier | Perceptual Speech |
| Imagine a person mumbling an apology | Perceptual Speech |
| imagine a tour guide speaking | Perceptual Speech |

| Prompt | Category |
|---|---|
| Imagine a waiter asking for an order | Perceptual Speech |
| Imagine a pilot on an airplane intercom | Perceptual Speech |
| Imagine a congested person speaking | Perceptual Speech |
| Imagine hearing a phone voice assistant | Perceptual Speech |
| Imagine ordering at a drive-thru | Perceptual Speech |
| Imagine a person speaking while chewing | Perceptual Speech |
| Imagine a hoarse person speaking | Perceptual Speech |
| Imagine a person telling a story | Perceptual Speech |
| Imagine listening to an audiobook | Perceptual Speech |
| Imagine an auctioneer speaking | Perceptual Speech |
| Imagine repeating your family's names | Inner Speech |
| Imagine words that start with B | Inner Speech |
| Imagine reciting famous movie titles | Inner Speech |
| Imagine words that rhyme with ice | Inner Speech |
| Imagine counting down from 20 | Inner Speech |
| Imagine the names of US presidents | Inner Speech |
| Imagine listing the US states | Inner Speech |
| Imagine listing names of fruits | Inner Speech |
| Imagine words that rhyme with pan | Inner Speech |
| Imagine counting up from 50 | Inner Speech |
| Imagine the names of schools you've attended | Inner Speech |
| Imagine explaining a recipe | Inner Speech |
| Imagine giving directions to your home | Inner Speech |
| Imagine the names of major cities | Inner Speech |
| Imagine words that rhyme with row | Inner Speech |
| Imagine your family's birth dates | Inner Speech |
| Imagine reciting the days of the week | Inner Speech |
| Imagine the names of body parts | Inner Speech |
| Imagine explaining your job | Inner Speech |
| Imagine reciting the months of the year | Inner Speech |
| Imagine listing genres of movies | Inner Speech |
| Imagine describing the weather | Inner Speech |
| Imagine words that start with T | Inner Speech |
| Imagine the words to Happy Birthday | Inner Speech |
| Imagine reciting your phone number | Inner Speech |
| Imagine reciting the alphabet from P | Inner Speech |
| Imagine counting up from 5 | Inner Speech |
| Imagine listing types of jewelry | Inner Speech |
| Imagine words that start with S | Inner Speech |
| Imagine the names of instruments | Inner Speech |
| Imagine giving an excuse | Inner Speech |
| Imagine counting down from 25 | Inner Speech |
| Imagine words that start with C | Inner Speech |
| Imagine listing names of countries | Inner Speech |
| Imagine words that rhyme with cat | Inner Speech |
| Imagine reciting colors of the rainbow | Inner Speech |
| Imagine listing names of animals | Inner Speech |
| Imagine listing types of clothing | Inner Speech |
| Imagine reciting the alphabet | Inner Speech |
| Imagine reciting your street address | Inner Speech |
| Imagine a trumpet playing 'Happy Birthday' | Sound |
| Imagine you can hear a babbling brook | Sound |
| Imagine the sounds of a construction site | Sound |
| Imagine elevator music | Sound |
| Imagine the sound of a dog barking | Sound |
| Imagine hearing a car engine start | Sound |
| Imagine the sound of someone typing | Sound |

| Prompt | Category |
| --- | --- |
| Imagine the sound of splashing water | Sound |
| Imagine the sound of birds singing | Sound |
| Imagine the sound of a fire engine | Sound |
| Imagine a violin playing a tune | Sound |
| Imagine a guitar being played | Sound |
| Imagine hearing chalk on a chalkboard | Sound |
| Imagine an opera singer's voice | Sound |
| Imagine the sound of crickets chirping | Sound |
| Imagine singing a karaoke song | Sound |
| Imagine hearing a vacuum cleaner | Sound |
| Imagine hearing a piece of paper being torn | Sound |
| Imagine the sound of popcorn popping | Sound |
| Imagine hearing a waterfall | Sound |
| Imagine the sound of a crowd applauding | Sound |
| Imagine the sound of a clock ticking | Sound |
| Imagine a police siren | Sound |
| Imagine the roar of a lion | Sound |
| Imagine a church organ playing | Sound |
| Imagine a doorbell ringing | Sound |
| Imagine hearing church bells tolling | Sound |
| Imagine hearing a phone ringing | Sound |
| Imagine the sound of wood being sawed | Sound |
| Imagine the sound of radio static | Sound |
| Imagine cars honking in city traffic | Sound |
| Imagine the sound of someone sneezing | Sound |
| Imagine hearing a choir singing | Sound |
| Imagine hearing a microwave beep | Sound |
| Imagine a saxophone playing | Sound |
| Imagine you can hear a helicopter | Sound |
| Imagine a baby crying | Sound |
| Imagine a racecar driving past | Sound |
| Imagine a piano being played | Sound |
| Imagine a rock song playing on the radio | Sound |
